# phyB photobodies display molecular memory, regulating light signaling transcriptional states through phase separation

**DOI:** 10.64898/2026.09.06.749697

**Authors:** Freddy Igiebor, Laeschkir Würthner, Franziska Stamm, Subiya Haque, Ondrej Havlicek, Janina Schmidt, Merve Cetiner, Ivan Zubcic, Omar Heliel, Lauren M Saunders, Isabella R. Graf, Kasper van Gelderen

## Abstract

Phytochrome B is a crucial red and far-red light sensor, which controls plant development and environmental responses to light and temperature. In the nucleus, phyB forms phase separated condensates, also called photobodies, consisting of various transcriptional regulators and transcription factors. phyB photobodies can form or disperse depending on the amount of light, the light quality, or ambient temperature. However, the biological relevance for forming photobodies is still unclear. In recent years the study of condensates has given rise to the idea that condensates can act as molecular memory, either due to long equilibration timescales arising from strong interactions that can kinetically trap a mixture in a phase-separated state beyond the equilibrium coexistence line, or from phase-separation dependent modification processes or transcriptional regulation. Here, we hypothesize that the formation of phyB photobodies and the associated sequestration of signaling molecules allows for the formation of cell-type specific memory about light conditions. To address this hypothesis, we first developed a live imaging setup of phyB photobodies which shows that they do not follow classic liquid-liquid phase separation dynamics, but instead have a restriction on their size, intensity and minimum number. Fluorescence recovery after photobleaching (FRAP) then showed that photobodies between cell types have different levels of mobilities and diffusivities, suggesting that they are more stable in darkness or high red light, versus low red light. These differences in phyB photobody stability between cell types were also confirmed through whole-mount confocal imaging and single-nucleus transcriptomics in the same light-perturbation series. Through sci-Seq single nucleus spatial transcriptomics of the cotyledon in this light perturbation series, we identify a photobody-associated transcriptional cluster of nuclei that regulates photosynthesis, circadian rhythm, and RNA processing; the phyB photobody likely promotes its own stability by regulating *TZP* expression. The correlation between photobodies and transcriptional regulation shows how phyB condensates can provide cell-type specific light responses and exhibit signaling memory in changing environmental conditions.

## Introduction

Light is the most important resource for plant life, and to measure light quality and quantity, plants have evolved dedicated sensors. Phytochromes are a conserved family of bilin-binding photoreceptors found in plants, bacteria, and fungi. They primarily detect red (R) and far-red (FR) light and play fundamental roles in sensing the light and temperature environment to regulate growth and development across the entire plant life cycle, from seed germination to flowering (Legris et al. 2019; Cheng et al. 2021). Phytochrome B (phyB) is the most important red (600-700 nm) and far-red (700-750 nm) light sensor in plants, and is used to respond to changes in the light environment, mainly due to plant-plant competition and light quality (Ballaré and Pierik 2017; Hernando et al. 2021), but also temperature (Legris et al. 2019). phyB is a homodimer that absorbs red light through its phytochromobilin chromophore, which leads to a conformational change from the ground state (Pr) to the active form (Pfr). The Pfr state can absorb far-red light, which inactivates phyB and converts it to the Pr state (Li et al. 2011). Reversal to the ground state can also occur independently of light, and is accelerated at higher temperature (Jung et al. 2016; Legris et al. 2016; Klose et al. 2020). Within plant cells, phyB can be found in phase separated sub-nuclear bodies, called photobodies (Chen et al. 2003). These small (300 nm) bodies presumably act as centers of light signaling, and contain many light signaling molecules (Willige et al. 2024). phyB photobodies form through phase separation (Chen et al. 2022), and are therefore highly dynamic structures, which respond to red and far-red light (Van Buskirk et al. 2014) and temperature (Legris et al. 2016; Hahm et al. 2020). phyB photobodies have been shown to recruit PHYTOCHROME INTERACTING FACTORS (PIFs), which are transcription factors that regulate light responses downstream of phytochromes (Leivar et al. 2020). PIF recruitment to photobodies has been shown to cover different molecular functions. In general PIFs interact with active (Pfr) phyB, which leads to PIF inhibition and degradation (Ni et al. 2013, 2014). However, PIF3, PIF5 and PIF7 have also been shown to be stabilized in photobodies, with PIF7 being relatively stable and inhibited by phyB through association (Kim et al. 2023, 2024; Xie et al. 2023). Interestingly, in pull-down or proximity labelling mass spectrometry studies of phyB, PIFs have never been identified as key photobody components (Huang et al. 2016; Kim et al. 2023; Olson et al. 2025). This is most likely because of the transient nature of the phyB PIF interaction and low PIF stability. Finally, phyB photobodies are involved in regulating the spliceosome (Xin et al. 2017; Kathare et al. 2022; Yan et al. 2022), and they can be associated with particular chromocenters (Du et al. 2024).

Taken together, it is very likely that the function of the phyB photobody is to regulate transcription, or transcription-related processes, either by inhibiting or stabilizing transcription factors, or regulating splicing. However, it is unclear what those processes, PIF stability regulation, splicing, and transcriptional regulation could gain by a phase separated condensate state. Therefore, the question arises what the particular advantage of phase separation is for phyB.

Biological (liquid-liquid) phase separation has been studied as a biophysical phenomenon for more than 15 years, and is the process whereby proteins separate as a membraneless organelle from within a mixture of proteins, metabolites and nucleic acids (Hyman et al. 2014). Phase separation is a reversible, dynamic process, whereby there is continuous molecular interchange between compartments, which are stabilized by intermolecular interactions (Shin and Brangwynne 2017). Phase separated bodies (or droplets) are often promoted by the multivalent interactions from intrinsically disordered proteins, and other biomolecules such as RNA or ATP (Banani et al. 2017; Shao et al. 2022; Kota et al. 2024; Lu et al. 2024). Intrinsically disordered proteins (not to be confused with prions) lack a stably folded structure, but can conform dynamically, depending on the (protein) environment they reside in. PhyB photobodies also contain a multitude of intrinsically disordered proteins: PCH1 and PCHL (Enderle et al. 2017; Huang et al. 2019), ELF3 (Jung et al. 2020; Kim et al. 2023), TZP (Fang et al. 2022), and PIF7 (Xie et al. 2023), to name the most well-described examples. It has been shown empirically, and through mathematical modelling, that there are different degrees of persistence of phyB photobodies with larger photobodies in high light existing longer than those in low light, where photobodies are generally smaller (Buskirk et al. 2014; Trupkin et al. 2014; Klose et al. 2015a). PhyB photobodies can persist in solution (Kim et al. 2023), and recover after photobleaching (Chen et al. 2022; Kim et al. 2024). We hypothesize that the difference in stability between so-called ‘early’ and ‘late’ photobodies can be attributed to so-called coarsening dynamics (Hyman et al. 2014), where after nucleation of condensates, the number of condensates gradually decreases while the average volume of individual condensates increases. In equilibrium systems, one particular process leading to coarsening is called Ostwald ripening (Ratke and Voorhees 2002): over time small droplets shrink at the expense of larger ones in order to minimize the free energy of the system and thereby the total surface area between the dense and dilute phase. An idealized equilibrium coarsening process is described mathematically by the so-called LSW theory (Lifshitz and Slyozov 1961; Wagner 1961; Hyman et al. 2014; Shin and Brangwynne 2017; Berry et al. 2018), which also makes a specific prediction on how the average radius of droplets grows as a function of time.

Some instances of phase separation, including during thermosensing, have been proposed to act as a so-called ‘molecular memory’, in which a transient input signal leaves a long-lasting imprint on the localization and sequestration of associated signaling proteins and/or nucleic acids, for instance by phase-separation dependent transcriptional regulation or modifications. This memory mediated by phase separation then changes the response to a repeat of a stimulus, allowing for the control of signaling dynamics in response to environmental changes and developmental cues (Cuevas-Velazquez and Dinneny 2018; Dine et al. 2018; Murcia et al. 2021). In principle, molecular memory can also arise from hysteresis, where long equilibration timescales arising from strong interactions can kinetically trap a mixture in a phase-separated state beyond the equilibrium coexistence line (Garcia Quiroz et al. 2019). Molecular memory has already been observed in the temperature-dependent condensate formation of EARLY FLOWERING 3 (ELF3), a circadian and thermoregulatory intrinsically disordered protein that interacts with phyB (Murcia et al. 2022; Yang et al. 2025). Taken together, transient signals such as increased temperature or changes in the light condition, might leave a persistent imprint on phyB photobody formation and growth and thereby constitute a form of molecular memory.

In order to test how phyB photobodies grow and coarsen in response to light signals we first sought to investigate phyB photobody formation from first principles, and determine the biophysical parameters regulating phyB photobody phase separation. Interestingly, we found that phyB photobodies do not follow phase separation scaling laws as predicted by LSW Theory, but have a restriction on their size, intensity and minimum number. We then tested how much phyB photobodies can recover from photobleaching (FRAP) in different light conditions, and found that darkness leads to decreased mobility, while there are clear differences in FRAP rates between cell types. We used whole-mount imaging plus a custom deep learning image analysis pipeline to map cotyledon cell-specific phyB photobody responses to a light/dark treatment series and identified different response and coarsening behaviours for different cell types in response to the same light treatment. Furthermore, there were clear differences between the persistence of light states into the darkness in different cotyledon cell types. Finally, we performed a single nucleus transcriptomics experiment in parallel with the cell-specific imaging to correlate photobody behaviour with transcriptional output, where we identify a specific photobody-associated cluster of transcriptional regulation, which clearly links to photosynthesis, circadian rhythm, and RNA modification, and suggests a positive feedback loop to promote photobody stability, linking phyB photobody coarsening to light-signaling memory function.

## Results

### Time-course live imaging of phyB photobodies reveals that they do not behave as classical phase-separated liquids

PhyB photobodies have been recognized to form so-called ‘early’ and ‘late’ photobodies depending on the length of red light exposure (Meyer 2020). However, photobody formation and coarsening has never been characterized before in a time-resolved manner. We exposed five-day old cotyledons of *phyB-9 pphyB:phyB-YFP* seedlings (Viczián et al. 2020) to an hour of far-red light (730 nm, 30 µmol/m^2^/s) to completely abolish phyB photobodies in the pavement cells. We then used a set-up with a flexible LED light, illuminating the sample at the confocal microscope with 15 µmol/m^2^/s 670 nm red light and imaged pavement cells during a 2-hour time-course experiment with 5-minute intervals **(Figure 1A)**. To quantify the images we used a Biom3D image analysis deep learning model (Mougeot et al. 2026), which we trained using ∼100 annotated single confocal slices. We wrote a custom python script to pre-process the time-course data for Biom3D, and post process the data for analysis in R. We then compared our Biom3D method with a thresholding+clustering algorithm of ICY (HK-means) and hand counting, and found that Biom3D could annotate images better than HK-means, especially in low-contrast images, and was within 5% of hand counting results (**Figure S1, Supplemental methods**). Using this pipeline, we quantified photobody parameters of individual nuclei **(Figure 1B-E)**. Average photobody number per nucleus quickly decreased and stabilized over the course of 30 minutes and reached a stable level at around 20 on average **(Figure 1B)**. The average photobody volume per nucleus, average intensity and sum intensity increased sharply in the first 30 minutes and then plateaued **(Figure 1C-E)**. At first glance, this behaviour resembles the nucleation and growth behaviour of phase separation (Berry et al. 2018), and we fitted all the results with a single exponential curve each (orange line, Figure 1B), and compared whether these curves conform to LSW coarsening theory **(Figure 2)**. LSW theory states that: 1. The droplet number decreases in inverse proportion to time (N(t)∝t^(-1)), which on a log–log plot corresponds to a slope of −1. 2. The mean radius grows <R> ∼ t^(1/3), which then implies that the volume grows V∼<R>^3 ∼ t. 3. In our timelapse series, the radius and volume increased slower than expected **(Figure 2A,B).** The droplet number decreased slower than expected by LSW coarsening theory and furthermore reached a minimum number >1 of photobodies, which did not decrease over the course of this timelapse **(Figure2 C)**. So, once photobodies are nucleated and grown, they merge into larger condensates much less frequently than expected from the idealized Ostwald ripening. Instead of fusing continuously, they tend to remain as separate entities. We also imaged phyB photobodies using Airyscan super-resolution mode. Here we could see many smaller, low-intensity photobodies than we can see in normal confocal mode, even in a coarsened state (**Figure S2**). These observations strengthen the notion that phyB photobodies are not purely diffusion-limited condensates, where the molecular mobility is altered by addition of (de)stabilizing factors, or by biological structures in the nucleus such as chromatin. Finally, the total photobody material increased **(Figure 2D)**, which might be explained by nuclear import of phyB in red light (Klose et al. 2015b). Together, these findings indicate that diffusion-limited kinetics alone cannot explain photobody behaviour. In order to characterize the biophysical properties of phyB photobodies further and to elucidate how they differ between different cell types and change as a function of light stimulation, we determined interfacial molecular mobilities via Fluorescence Recovery After Photobleaching (FRAP) experiments.

**Figure 1.**
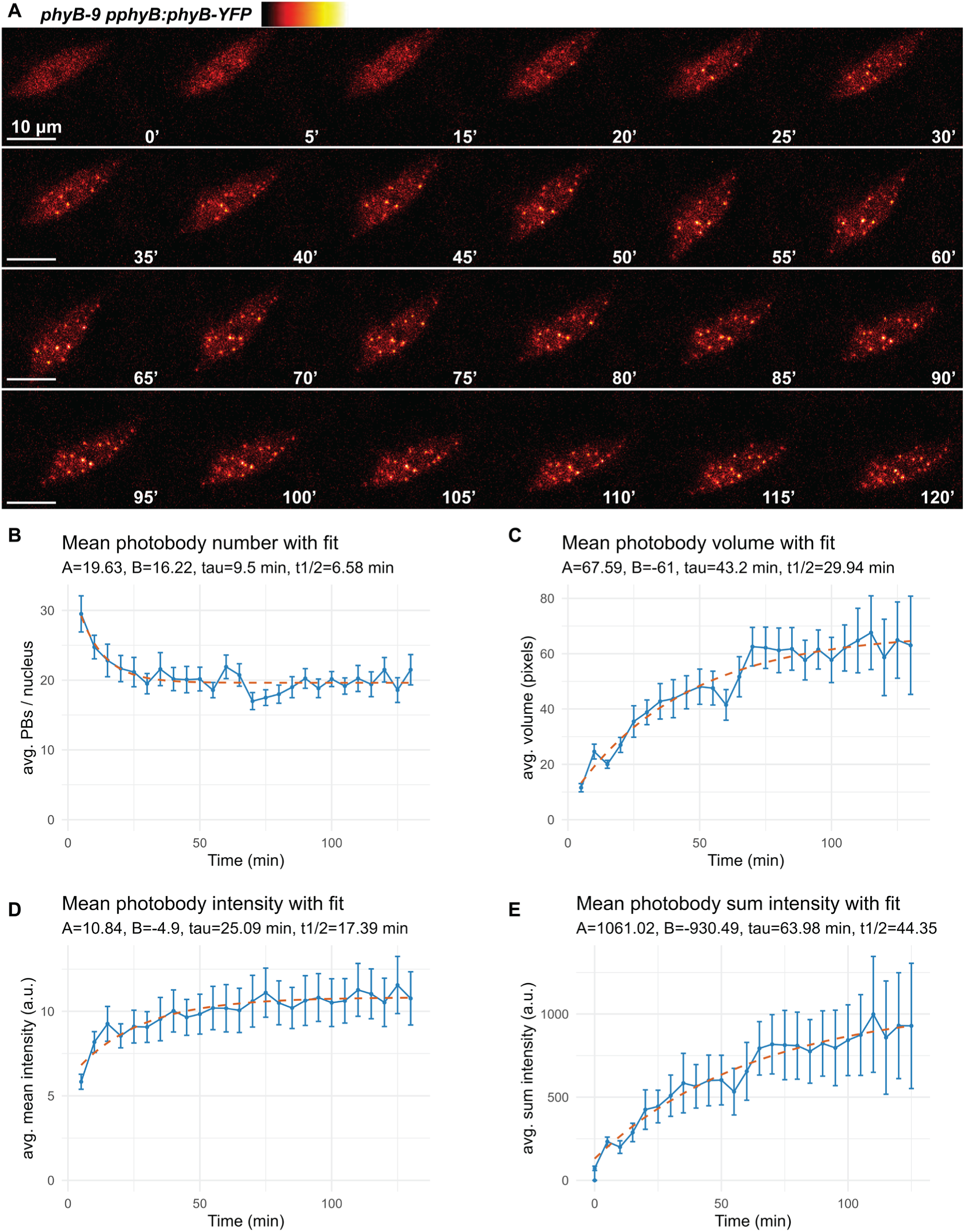
Time-course live imaging of phyB photobodies in pavement cells under red light shows coarsening behaviour. (A) Representative confocal images of a time-course experiment with pavement cell nuclei following transition from far-red light to continuous red light. Indicated are the minutes after far-red to red transition, scale bar = 10 µm. (B-E) Quantification of photobody parameters in the time-course experiment, displayed are the average values of 12 nuclei with a single-exponential curve fit (dashed line) showing parameter *A, B τ,* and calculated half-time, where *y = A +Be^-t/τ^* (displayed above graph). (B) Average number of photobodies per nucleus, (C) average photobody volume, (D) average intensity per photobody, (D) average sum intensity per photobody. Error bars display SEM.

**Figure 2.**
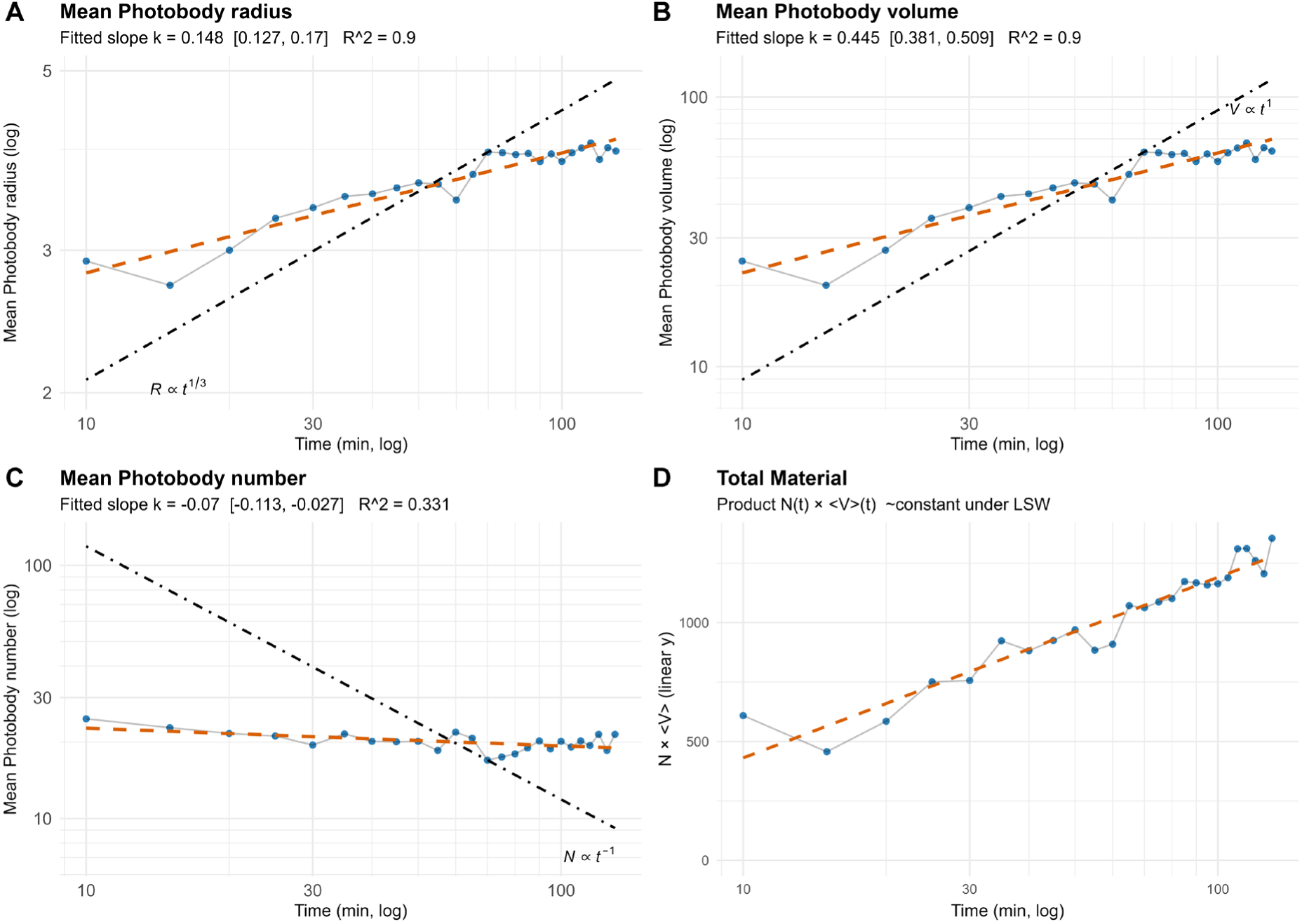
phyB photobody coarsening does not follow the power laws of Lifshitz–Slyozov– Wagner (LSW) theory. Plots showing the scaling relationships of the data from the experiment in Figure 1 in log scale (A-C), or linear (D), along with a fitted slope (orange dashed line) and the slope rule predicted by LSW theory (black dashed line). (A) Mean photobody radius: The fitted slope k ≈ 0.148; LSW-predicted scaling law (R(t) ∝ t^1/3). (B) Mean photobody volume: The fitted slope k ≈ 0.445; LSW prediction (⟨V(t)⟩ ∝ t^1). (C) Photobody number (per nucleus): The fitted slope was k ≈ −0.07; LSW prediction (N(t) ∝ t^−1). (D) Total material (sum intensity): LSW predicts this to remain constant, however the product of droplet number and mean volume (N(t) × ⟨V⟩) doesn’t remain constant, indicating potential flux or change in nucleus volume.

### FRAP Analysis of phyB photobodies shows light and cell-specific differences in photobody interfacial mobility

An important attribute of phase-separated condensates is that there is dynamic molecular exchange between the condensates and the cyto/nucleoplasm. It has been shown that phyB photobodies display FRAP recovery at a rate between 25% and 50%, which decreases when the amount of phyB protein increases (Chen et al. 2022; Kim et al. 2024). Furthermore, we know that there are red light-specific interactions of phyB with factors that stabilize phyB photobodies, such as TZP (Enderle et al. 2017; Murcia et al. 2021; Fang et al. 2022), and PCH1. In parallel, higher light, and thus higher phyB Pfr is associated with larger photobodies (Su and Lagarias 2007; Trupkin et al. 2014; Legris et al. 2016; Viczián et al. 2020). Different light intensities (10 versus 0.5 µmol/m^2^/s) can increase FRAP rates in a phyB overexpressing line (Kim et al. 2024). Protein overexpression can influence phase separation in a great deal (Hyman et al. 2014; Alberti et al. 2019), therefore we wanted to test FRAP rates in a native promoter complementing line of phyB. We performed FRAP experiments using the *phyB-9 pphyB:phyB-YFP* line under different light conditions (darkness, low red, high red, and white light). We utilized guard cells because of the relatively low mobility of their nuclei and photobodies. FRAP measurements were made of photobodies after 2h of darkness, 1.5 h high red light (60 µmol), 1.5 h low red light (15 µmol), and 100 µmol white light **(Figure 3)**. The FRAP data and subsequent curves were normalized so that the pre-bleach intensity was set at 100% and the post-bleach intensity at 0%. Then, FRAP data were corrected for photobleaching occurring through the post-bleach timecourse, using a control phyB signal in the image. A photobleaching experiment relies on a supply of non-bleached material being present to move/diffuse to the bleached area, thus we tested whether bleaching a guard cell with only one photobody would lead to a different outcome versus a nucleus with two or more, versus bleaching the whole nucleus. This showed clearly that bleaching a whole nucleus or a nucleus with one photobody recovered much less than a nucleus with two or more **(Figure S3)**, thus we proceeded to use guard cells with two nuclei or more. When comparing the different treatments, we observed that the FRAP recovery of guard cell nuclei of dark-treated cotyledons was lower and the recovery half time was slower than those in white light-treated cotyledons **(Figure 3A-C)**. High-red light treatment led to a faster and higher FRAP recovery than darkness, but lower than low red and white light pre-treatment **(Figure 3A-C)**. The maximum recovery rate is a metric of the mobility of molecules between the FRAP-ed and other compartments, thus dark-treated and high-red pre-treatment have less mobility between photobodies compared to low red light. The white light treatment had a red light component of equal intensity to the high red light, but was more similar to the low red light treatment, which may due to the additional white light wavelengths having additional / non-specific effects. Overall, these results clearly show that there are light-specific mobility differences between photobodies in guard cells.

**Figure 3.**
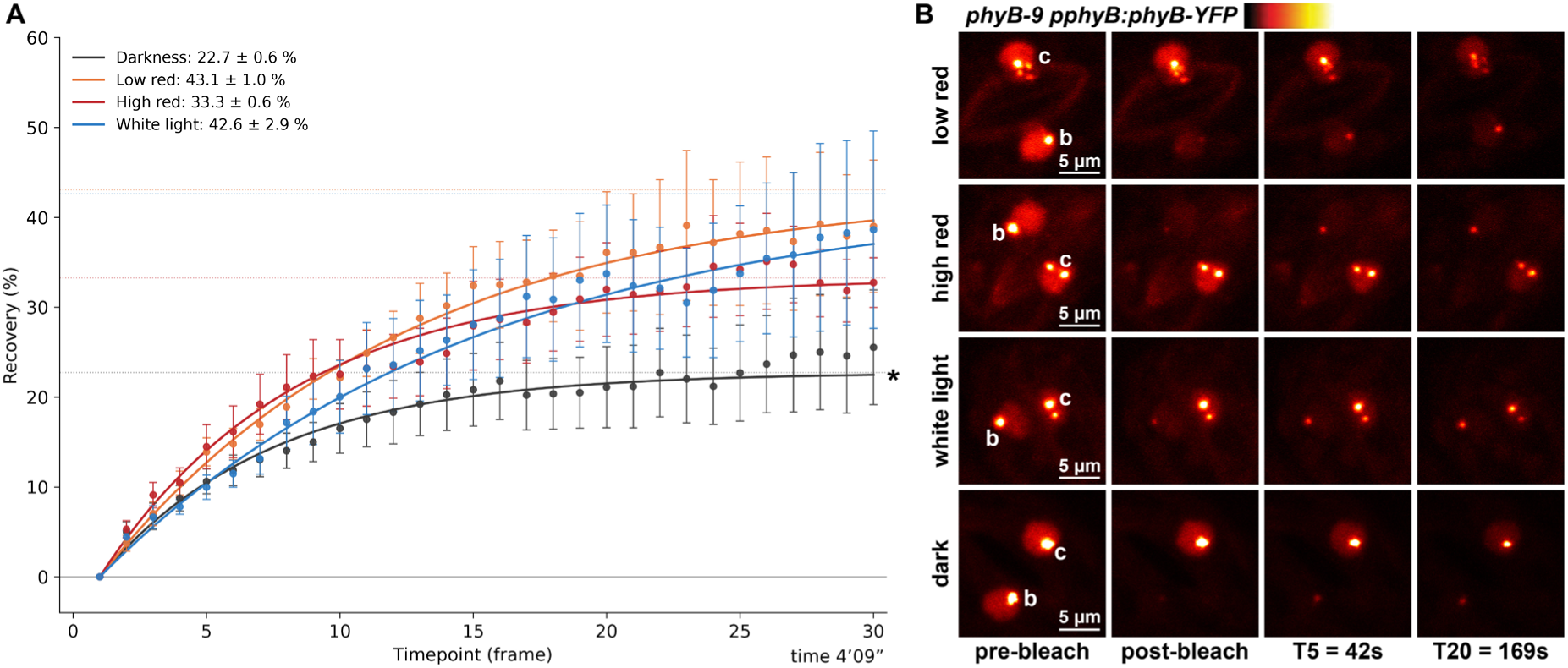
phyB photobodies have light-treatment FRAP recovery dynamics. Fluorescence Recovery After Photobleaching (FRAP) experiments using the *phyB-9 pphyB:phyB-YFP* line. (A) Guard cell FRAP recovery in four pre-treatment conditions: 2h of darkness, 1.5 h high red light, 1.5 h low red light, and constant white light. Photobody FRAP was monitored for 30 frames (8.47 seconds per frame, total duration ≈ 4 min 9 s), normalized to pre- and post-bleach, and corrected for general photobleaching. Displayed are the averages per time point with SEM (n=10), and a curve is fitted based on the average data, with the mobile fraction displayed in the top left legend, i.e., the percentage of fluorescence that recovered after bleaching based on the fitted curve. (B) Representative FRAP images from the dataset, with both guard cell nuclei shown. (b = bleached photobody, c = control photobody).

To obtain more relevant data for the cotyledon as a whole, we also performed FRAP in pavement cells. Again, we used the *phyB-9 pphyB:phyB-YFP* line, now under high (60 µmol/m^2^/s) and low (15 µmol/m^2^/s) red light conditions (660 nm), pre-treated by 1.5 hr darkness. In these scenarios, pavement cells have highly mobile photobodies (within the nucleus) and to be able to analyse them, we developed a 3D FRAP analysis method, dynaFRAP, which first uses Biom3D to segment all the photobodies in the timeseries and then tracks the recovering photobody during the analysis in the z-stack, applying normalization and photobleaching background correction **(see supplemental methods)**. Using dynaFRAP, we observed an increased maximum fitted recovery (expressed as the mobile fraction), in high red of 86%, while that of low red was 64% **(Figure 4A,B, Table S1)**. The recovery half time of high light was also faster (5.29 frames), than that of low red (6.77 frames). Furthermore, pavement cells in high red have a higher photobody number and photobody intensity compared to low red **(Figure S4)**. This shows that photobodies in pavement cells in high red light have more mobility between bodies, than those in low red light. In a separate experiment, analysed with dynaFRAP, guard cells also had a faster FRAP recovery half time in high red light (6.06 frames vs 11.35 frames), but no difference in mobile fraction, which was 33% and 31% (**Figure S5, Table S1**). This indicates that there are large differences between FRAP rates in different cell types. Furthermore, the FRAP recovery measured with the native promoter phyB-YFP line is significantly higher than what was measured before with phyB overexpression lines (25-50% (Kim et al. 2024), indicating the effect of raised expression levels on FRAP and showing that there is very high molecular interchange happening in phyB photobodies at native expression levels.

**Figure 4.**
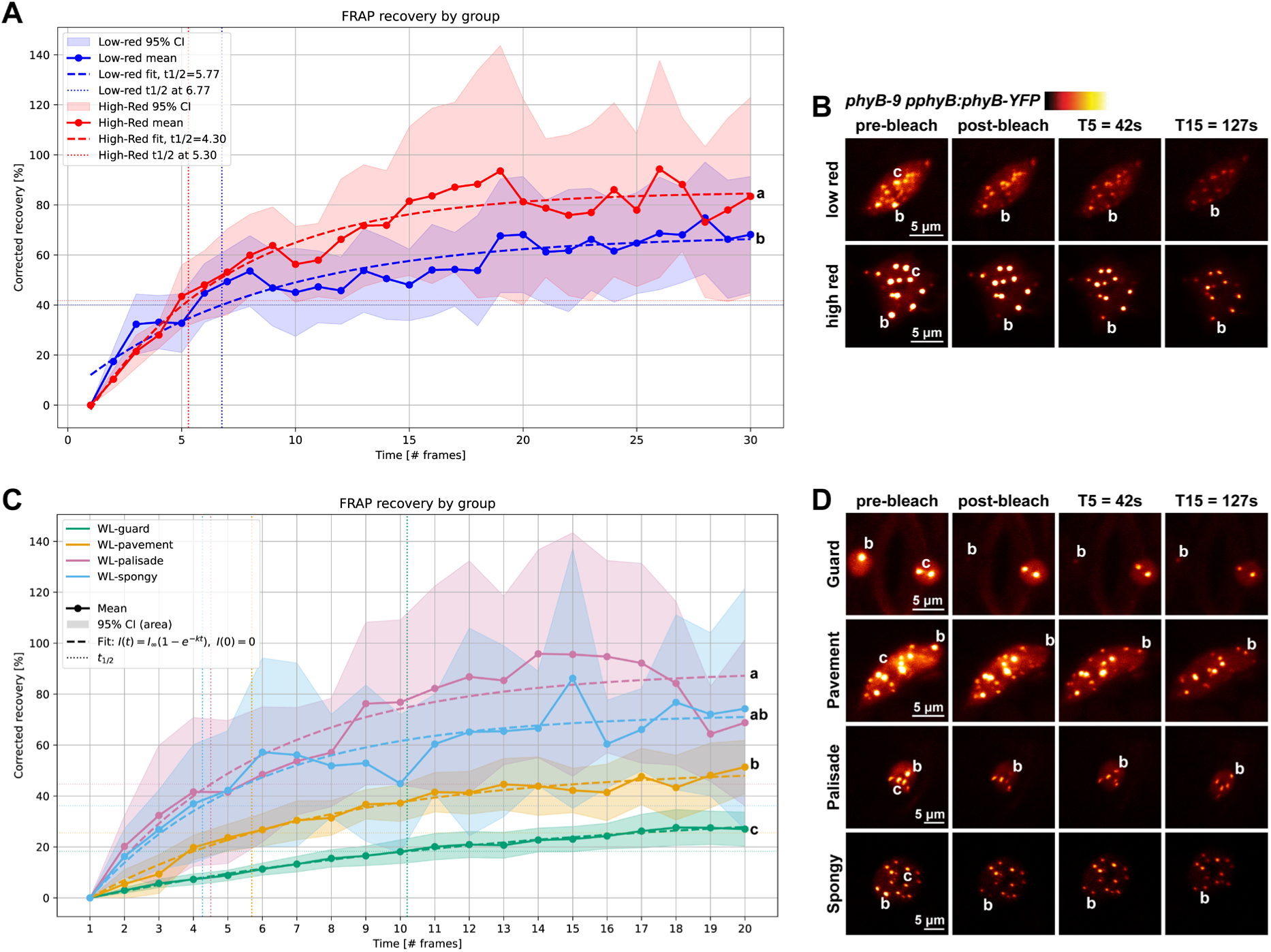
phyB photobodies have light-treatment and tissue-specific FRAP recovery dynamics. Fluorescence Recovery After Photobleaching (FRAP) experiments using the *phyB-9 pphyB:phyB-YFP* line. (A) Pavement cell FRAP experiment in red light (660 nm), at low (15 µmol/m^2^/s) or high (60 µmol/m^2^/s) intensity. Mean normalized FRAP recovery rates (shaded areas show 95% interval) with exponential fitted curves for photobody FRAP recovery. (B) representative images of experiment in (A). (C) FRAP recovery curves in white light pre-treatment of Guard, Pavement, Palisade and Spongy cells, similar to (A). Letters at end of curve denote statistically significant groups according to the curve plateau (One-way ANOVA, Holm pairwise test, p<0.05). (D) Representative images of experiment in (C).

To delve further into the cell-specific photobody differences in mobility, we performed FRAP on white-light grown seedlings comparing guard, pavement epidermis, and spongy mesophyll cells, using dynaFRAP analysis **(Figure 4).** Mesophyll cells in general had the highest mobile fraction (Palisade 87%, Spongy mesophyll 79%), which was significantly higher than pavement cell photobodies (56%), which was higher than that of guard cells (32%, **Figure 4C,D, Table S1)**. The recovery half time of the mesophyll was also fastest (Palisade 4.35 frames, spongy 5.13 frames), followed by pavement (6.89 frames) and guard cells (8.53 frames, **Figure 4C,D**. These results show clearly that there are cell-specific differences in photobody FRAP rates. Thus, we asked the question: do photobodies in different cells react differently to light and light perturbations?

### Whole-cotyledon imaging of phyB photobodies reveals distinct condition and cell-type specific changes in photobody stability

The bulk of phyB photobody data originates from the cotyledon pavement or hypocotyl epidermis cells (Chen et al. 2003; Su and Lagarias 2007; Buskirk et al. 2014; Legris et al. 2016; Hahm et al. 2020). These are specialized, endoreduplicated cells that are not necessarily representative for the response of the plant, organs, or tissues underneath (Tsukaya 2019). Using fixation and a clearsee clearing protocol (Ursache et al. 2018; Kurihara et al. 2021), we mapped phyB photobody responses to different durations of far-red light in the guard cells, pavement epidermis, palisade mesophyll and spongy mesophyll cells **(Fig. 5)**. This revealed that each cell type responds differently to the far-red light stimulus. Specifically, the epidermis cell photobodies disappeared after 10 minutes of far-red light, while even 30 minutes of far-red was not enough to dissolve all photobodies in the guard cells and spongy and palisade mesophyll **(Fig. 5A)**. Pavement cell photobodies fully disappeared and guard cell photobodies had an intensity, but no size decrease, while palisade and spongy mesophyll had both a size and intensity decrease **(Fig. 5B)**. After one hour of far-red light all photobodies fully disappeared in all cell types.

**Figure 5:**
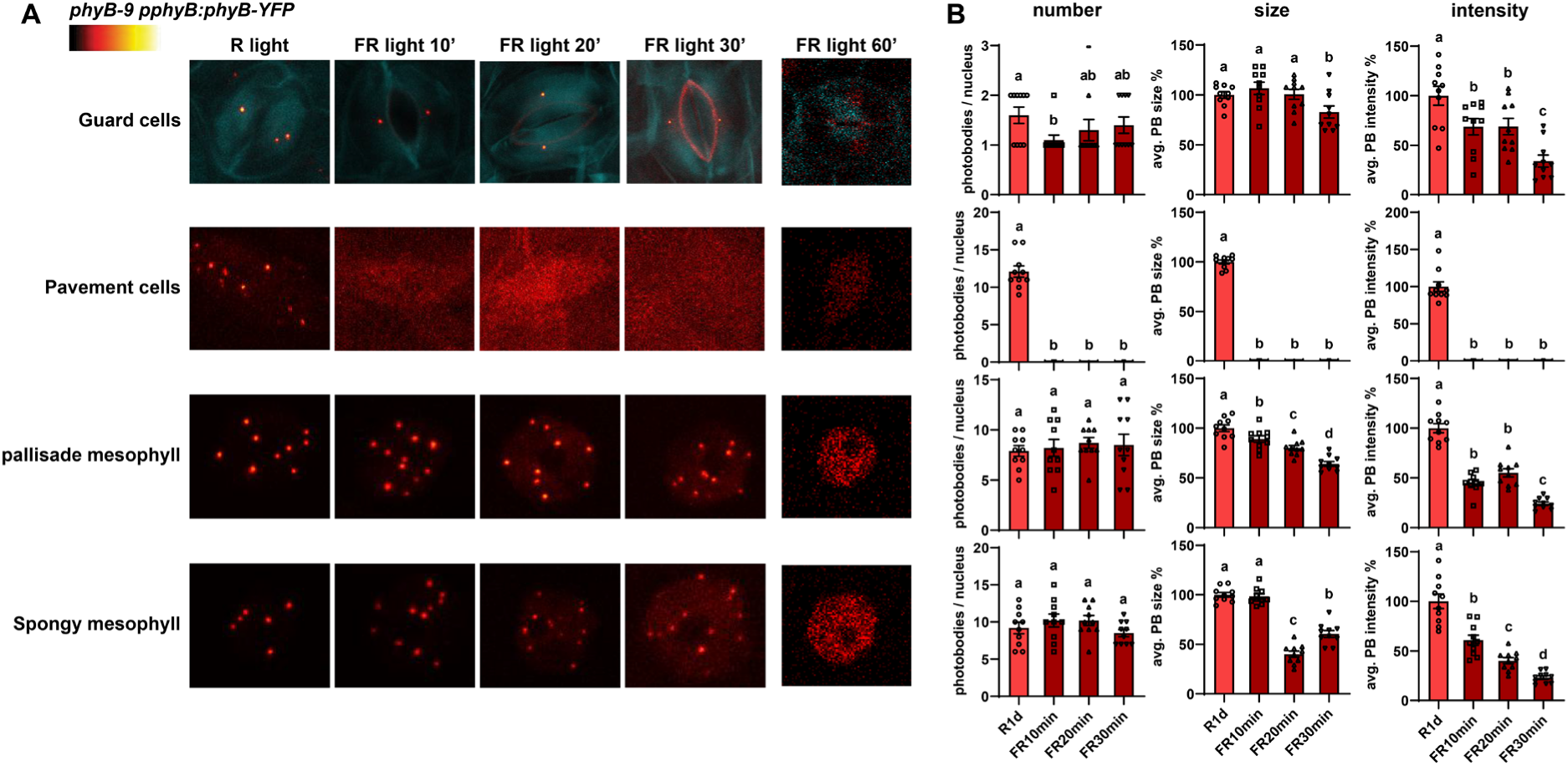
Pavement cells respond faster to far-red light than mesophyll cells. (A-D) Fluorescence images of phyB-YFP photobodies in different tissues of cotyledon. Seedlings were grown in white light for 4 days, followed by their respective light condition (R: red light 660 nm, 30 µmol/m^2^/s; FR: far-red light 730 nm 30 µmol/m^2^/s). (A) The membrane was dyed using DirectRed23. Scale bare represents 10 µm. Panel right: (A-D) Presented are the number, surface and average intensity of photobodies of individual tissues of the cotyledon. Photobodies of ten nuclei per layer of one cotyledon were taken into account. Seedlings were grown in white light for 4 days, followed by their respective light condition (R: red light; FR: far-red light). Error bar represents minimum to maximum values.

This simple experiment showed that there is a differential response between cell types to far-red light. If we assume that phyB molecules have the same photochemistry in each cell, then there need to be other factors, such as expression level, or stabilizing interactors that cause a difference in persistence of photobodies. Through these mechanisms, phyB photobodies can potentially retain memory about past light stimulation: by persistent changes following a light condition and thereby reacting to a repeated stimulus in a progressively changed manner. To test the idea that phyB photobodies display conditional-specific memory, we performed a time-series experiment starting with a 12h darkness treatment (simulating the night), then followed by a 1.5hr red light treatment (start of day), interrupted by a 4h dark treatment, (simulating sudden deep shading, submergence, or similar), followed by reappearance of red light for 20 minutes. This setup allows us to observe how photobodies react to a repeated stimulus and how they retain memory about past stimuli. We imaged whole-mount fixed, clearsee cleared, cotyledon samples of phyB-YFP at very high resolution with Airyscan. We manually sorted cell types based on cell and nuclear shape and location within the z-stack (guard, pavement, palisade mesophyll and spongy mesophyll) **(Figure 6A)**, and used our Biom3D pipeline to analyze the photobodies. We then plotted the volume and intensity of each photobody per cell type in a scatterplot, and the photobody number/nucleus in a boxplot **(Figure 6B,C** all plots: **Figure S5).** Strikingly, we found that that after 12 hrs of darkness, bright photobodies within the mesophyll and guard cells still persist, while in the pavement cells, only faint, small photobodies are left (**Figure 6A,B**), showcasing a differential response of phyB photobodies in different cell types. In all cell types we see a direct response to 5 and 20 minutes of red light, in that smaller and low intensity photobodies are formed. These quickly coarsen over the time period of 1.5 hrs, which occurs faster in the pavement cells than any other cell type, as evident by the quick reduction in photobody number (**Figure 6C**) and the increase in volume and intensity **(Figure 6B**). The guard, spongy and palisade mesophyll cells coarsening process continues throughout the second darkness phase, since photobody number is still decreasing and intensity and size are increasing until 2 to 3 hrs darkness treatment **(Figure 6B,C Figure S6**), again indicating a persistence of the light state. Applying darkness for 1 hr after red light elicits an immediate reaction from the pavement cells, but not so in the mesophyll cells. After 4 hrs darkness, the photobodies in the guard cells have reduced to one and those in the pavement cells have reduced to a situation similar to the 12h dark treatment **(Figure 6 B,C Figure S6**). At 4h darkness the photobodies in the spongy mesophyll and palisade cells also have stabilized to a situation similar to the previous 12h dark treatment, except for the absolute number of photobodies per nucleus, which was lower than after 12h darkness **(Figure 6B,C)**. Reapplication of red light led to a photobody response in all cell types, however, this response was different from first 20-minute light application. Photobody number, intensity and volume for palisade and spongy mesophyll cells and the pavement cells differed from the first 20-minute light application **(Figure 6A-C**). This is a clear example of a memory state of the phyB photobodies. Overall, this light series shows that the pavement cells react most quickly and flexibly to changing light environments. The other three cell types respond less, with the spongy and palisade mesophyll showing photobody persistence after 12hrs darkness. Similarly, we saw that these cell types are also resistant to far-red light. Furthermore, the coarsening process in the mesophyll continued in the absence of light. Thus, photobodies retain memory about light stimulation, in the face of phyB inactivation, with cell-specific differences. Next, we wanted to know if these cell- and light-specific photobody changes can be correlated to changes in transcriptional outputs. Thus, we performed a single-nucleus sequencing experiment in the same light perturbation series as **figures 5 and 6**.

**Figure 6.**
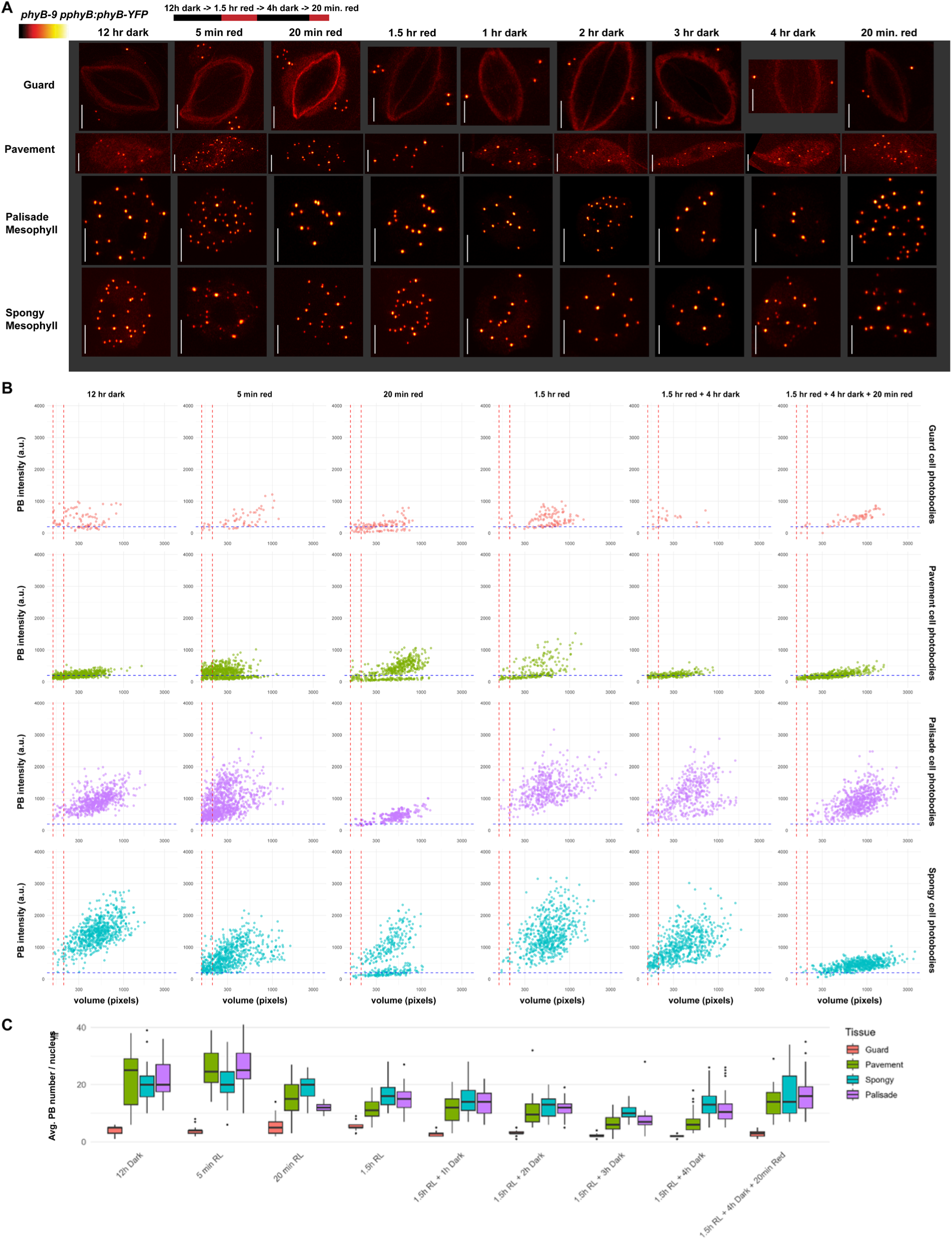
Mesophyll and pavement cells conserve either light, or dark information, and display memory. (A) Representative images of a Confocal Airyscan microscopy experiment of individual nuclei across four tissue types (Guard, Pavement, Spongy, and Palisade), and selected consecutive light and dark treatments (12 h Dark -> 1.5 h Red Light -> 4 h Dark -> 20 min Red Light), fixed and clearsee-cleared at different indicated timepoints during the series (Scale bar = 5 um). (B) Quantified data of photobody intensity and volume taken from the dataset (A), plotting individual for volume and intensity. Each point represents a single annotated photobody. The dashed red vertical line indicates the applied threshold for photobody intensity, while the dashed blue horizontal line indicates the threshold for photobody volume. (C) Photobody number / nucleus of data in (A,B). A two-way ANOVA showed significant effects of treatment and tissue type (p < 0.001) for volume, intensity and photobody number.

### Sci-seq single-nucleus RNA sequencing identifies photobody-response-specific cluster

To investigate the transcriptional responses associated with our light perturbation series in a cell-specific manner we used single nucleus combinatorial indexing-based RNA-sequencing (sci-RNA-Seq) (Martin et al. 2023). We sampled cotyledons of Col-0 and selected mutants in response to the same light perturbation setup as the cell-specific photobody imaging series, taking four timepoints right before each change in light conditions **(Figure 7B)**.

**Figure 7.**
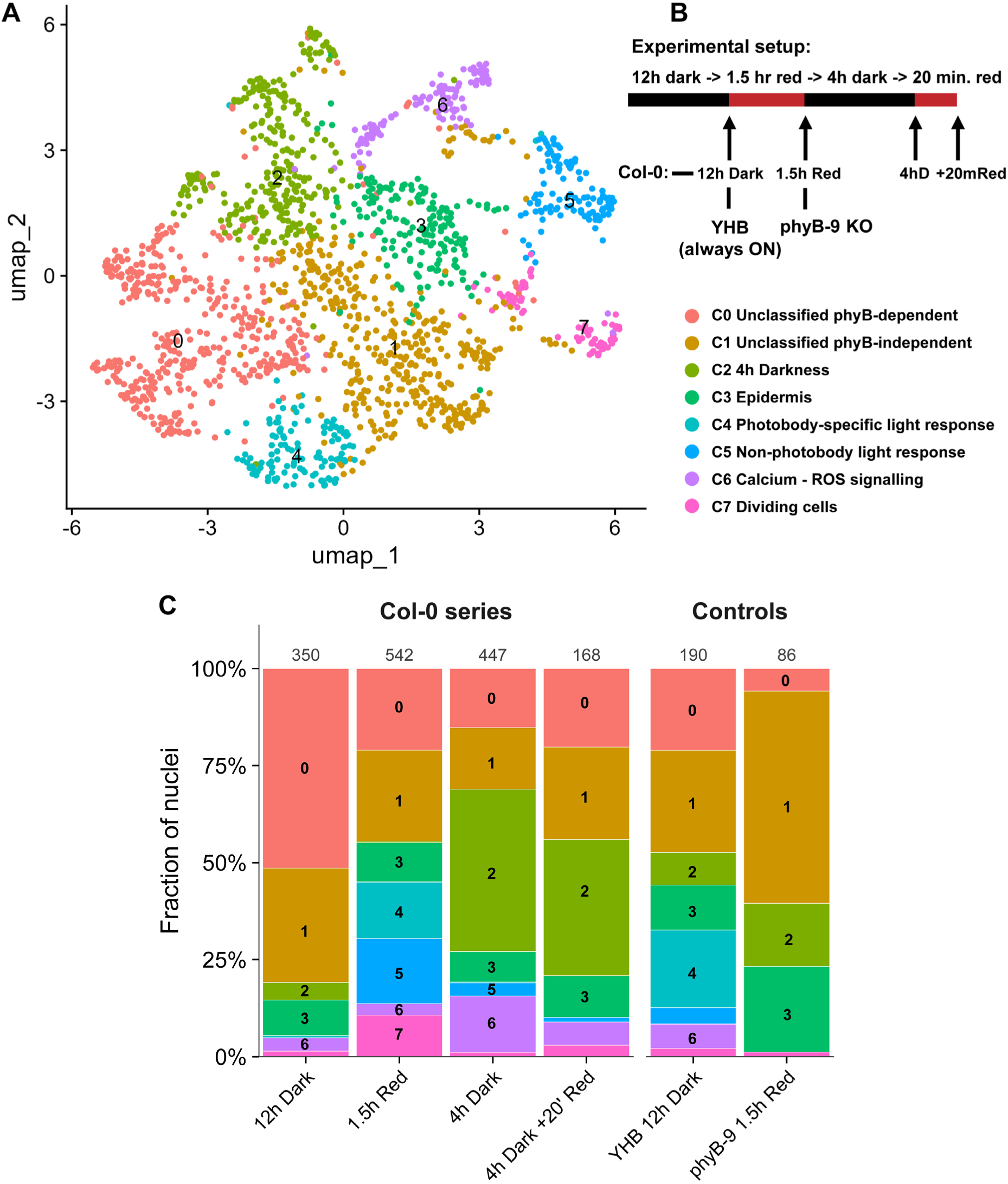
Single Nucleus RNA-seq shows light- and cell-type specific responses. A single-nucleus RNA-seq experiment was performed according to the scheme on the right of the panel (B), similarly to figure 5. (A): UMAP plot showing expression clusters of individual nuclear transcriptomes, with classifications of the clusters based on GO term enrichment of the cluster-specific marker genes (893 DEGs). (C) Barplots of cluster distribution per treatment from (A), showing the contribution of each treatment type within the experiment. For individual UMAP plots, **see Figure S11**)

We used the *35S:phyB^Y267H^-YFP (YHB)* line in darkness as a positive control for photobody-related signals, since YHB is a variant of phyB that is always in the Pfr conformation, independent of the light conditions, and thus always produces photobodies (Su and Lagarias 2007). Conversely, we used *phyB-9* in red light as a negative control against phyB-related signals. After quality control of our expression data **(Figure S7),** we identified 8 main clusters in our dataset (**Figure 7A)**. Using a published dataset (Lee et al. 2025) to assign cell types we were able to confidently call Mesophyll and Epidermis cells, and with less confidence the difference between Spongy and Palisade mesophyll **(Figure S8)**. Overall the spongy and palisade mesophyll had significant overlap and thus we grouped them together. Vascular tissue and guard cells were detectable but too low in abundance. Cluster 3 was clearly identified as the epidermal (pavement) cell type **(Figure 7A),** however most of these clusters did not represent a particular cell type, but rather a cell state of a particular subset of mesophyll-enriched nuclei **(Figure S9A-D)**. We performed a GO-term analysis on the DEGs associated with each cluster **(Figure S10, Supplemental file 1)**. Cluster 2 had a clear darkness / senescence / catabolic signature and was highly enriched in the 4h Dark treatment **(Figure 7C, S11)**. Clusters 4 and 5 both were enriched in the 1.5h light treatment, and not present in *phyB-9*, but Cluster 4 was especially prevalent in *YHB* in the dark **(Figure 7C)**. Cluster 4 had the most DEGs (429) and had the clearest GO term enrichment of photosynthesis, light perception and circadian rhythm, and specifically, mRNA binding for molecular function GO **(Figure S10, Supplemental file 1)**. Particularly noteworthy is that DEGs defining cluster 4 include *phyB* itself; *PIF5, SPA1, SPA3, COP1 and TZP,* all of which are interactors with phyB and are upregulated; *LHY, CCA1, RVE8, LNK1, LNK3, RPT2* and *PHOT2,* which are important circadian regulators. Therefore, Cluster 4 is the cluster that best correlates with photobody behaviour and *phyB-*dependent light regulation **(Figure 7C, S11, Supplemental file 1)**. Of the DEGs in Cluster 4, 8 genes, including *PIF5, phyB, CAB4, LNK1, LNK3* and *PIN3*, are also found as phyB DNA binding sites at 17°C white light in a ChIP-seq experiment **(Table S2)** (Jung et al. 2016), further strengthening the notion that Cluster 4 is the photobody-related expression cluster. Cluster 5 on the other hand does not seem to have a clear phyB association, nor a clear GO term enrichment, and appears to be a more generic light response **(Figure 7C, S10, S11, Supplemental file 1)**. Cluster 6 was not enriched for a particular light or dark state, but is more associated with stress, calcium signalling and ROS signalling **(Supplemental file 1)**. Cluster 7 is representing dividing cells, since it has clear GO enrichments associated with cell division processes (**Figure S10)**. Clusters 0 and 1 did not have a clear classification based on cell type nor GO term, and most likely represent low depth nuclei or shared background processes in the mesophyll. However, Cluster 0 was absent from the *phyB-9* sample, while Cluster 1 was not, marking Cluster 0 *phyB* dependent **(Figure 7C, S11)**.

We compared the DEGs from our dataset against existing *Arabidopsis* transcriptomics dataset DEGs and identified a high overlap with red light / phytochrome related experiments (Sakuraba et al. 2014; Shikata et al. 2014), circadian rhythm (Covington et al. 2008; Rugnone et al. 2013; Blair et al. 2019; Qin et al. 2025), and dark starvation / senescence (Liu et al. 2017; Kim et al. 2018) **(Figure S12a)**. Not surprisingly, there was a significant overlap between our differentially expressed genes and previous single nucleus leaf datasets (Tenorio Berrío et al. 2022; Xia et al. 2022; Qin et al. 2025). Clusters 1,2,3, 4 and 7 accounted for most of the overlap, especially with light- and circadian regulated datasets. The experiment from Rugnone et al. 2013 was of particular interest, since this dataset contained a treatment not dissimilar to the one described in this article, giving a disrupting light treatment during a darkness period in the subjective night, and a light treatment during darkness in the subjective day (Rugnone et al. 2013). Cluster 4 had a particularly high overlap with highly light-induced genes in both the night light treatment and subjective day light treatment, thus cluster 4 did not show a preference for night- or day-gating **(Figure S12B)**. Meanwhile clusters 1 and 2 had a strong overlap with the light-repressed genes in Rugnone et al. 2013, also not in a night-, or day-gated manner **(Figure S12B)**. This showed, that although our dataset has a significant overlap with circadian and light-regulated datasets, that our experiment is mostly a light-specific response rather than a circadian response. that Cluster 4 is a light response and not only a circadian gated response.

### DEG gene expression pattern analysis shows epidermis regulating light signaling and growth, while pavement cells control photosynthesis

To gain more insight in the processes and changes occurring between the different light treatments in the pavement and mesophyll cells, we analysed the temporal expression patterns of the Col-0-specific DEGs in our sci-seq dataset and performed a clustering analysis across the treatments. We identified specific sets of DEGs with shared expression patterns (now called blocks) across light treatments in epidermis, mesophyll or both (**Figure 8, S13A** for GO terms**, Supplemental file 2** for individual genes). Firstly, the 12h Dark treatment, was associated with block 3, which was too small to be classified according to a GO term. After the first light treatment (1.5h Red), we could identify expression blocks 1&2, specific for this treatment, which together have a high overlap with the photobody-associated cluster 4 **(Figure 8, see Jaccard-test inset)**. Block 1 is specifically upregulated in the mesophyll and consists mainly of photosynthesis related genes, such as light harvesting complexes and chloroplast regulators **(Figure 8, S13A)**. Block 2 is upregulated in both the epidermis and mesophyll and is enriched in circadian rhythm and RNA processing genes **(Figure 8, S13B)**. Block 2 also contains most of the previously mentioned light and circadian regulator genes from the photobody-specific Cluster 4: *SPA1, SPA3, COP1, TZP, LHY, CCA1, RVE8, RPT2, PHOT2, and PIF4* (not *PIF5*). Blocks 4 and 5 are specific for the dark perturbation treatment (4h Dark), with block 4 more prominent in the mesophyll, and block 5 in the epidermis, and both having strong overlap with the 4h dark cluster 2 **(Figure 8, S13C,D)**. Both blocks are enriched for catabolic, and absence of light terms, such as the starch degradation protein SEX1 (block 4), and senescence-associated genes *DIN2, DIN4, DIN10, ATG8B* (block 5). Block 4 also contains genes regulating pathogen defense: *DMR6, BBD1, PICBP, RPP4, GRIK2*. Block 6 is the largest block, and is epidermis specific for the second light treatment (+20 min. Red). Interestingly, it is enriched in hormonal signaling, especially cytokinin, ABA, brassinosteroids and auxin signalling genes, and contains growth regulators, such as *ARF6, FER, KAN, bHLH61* and *bHLH121* **(Figure 8, S13E)**. Block 6 also contains the essential circadian regulators *PRR5, PRR7 and PRR9*, and is again enriched for RNA processing, mainly splicing factors. Blocks 7 and 8 are smaller and mesophyll specific for the 20 minute red light treatment, and represent photosynthesis *(PSBA, PSBC, PSBM,* Block 7), and growth regulation (mainly auxin and flowering, such as GI, block 8) (**Figure S13F,G**).

**Figure 8.**
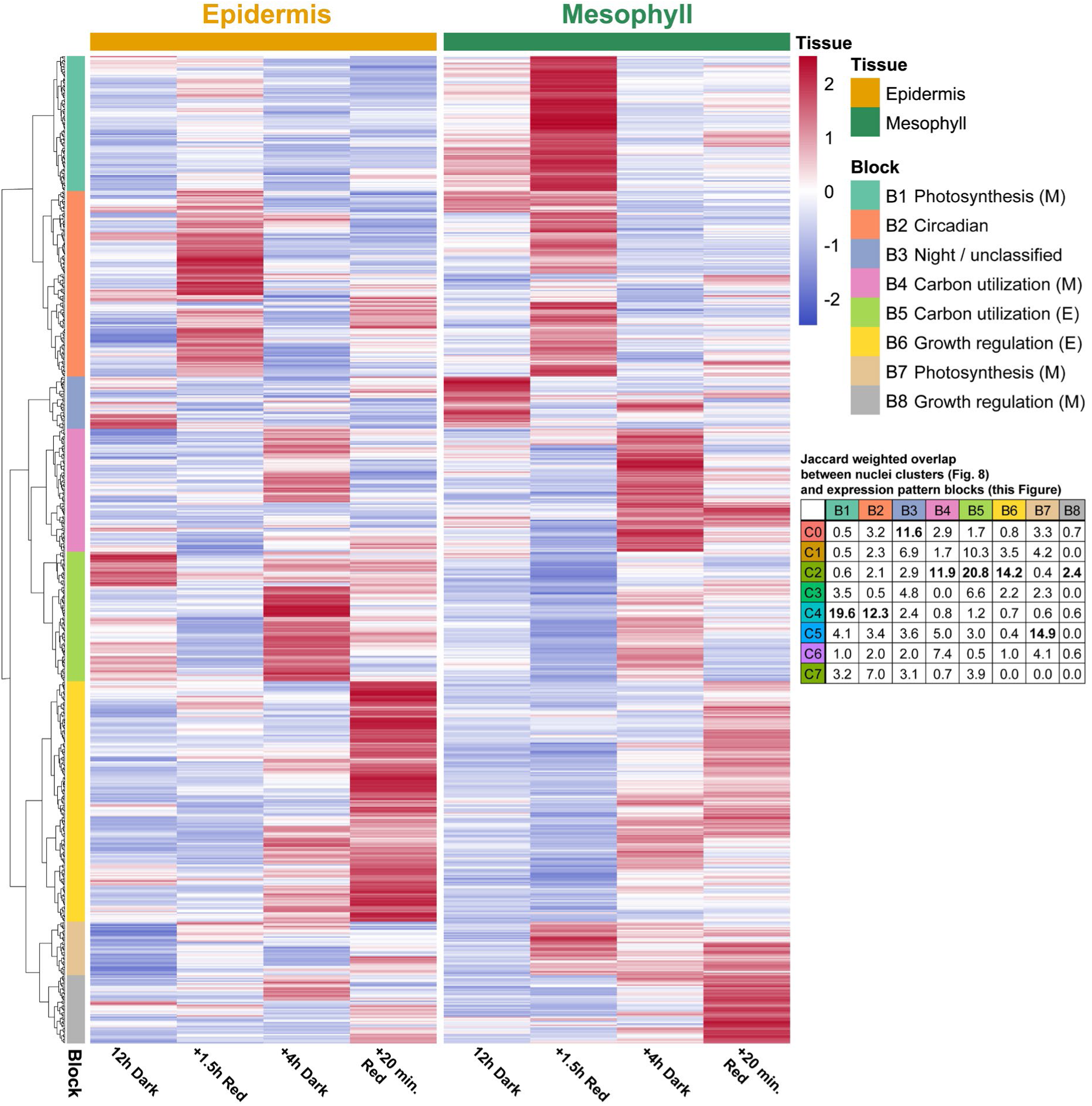
DEG expression pattern analysis shows how the mesophyll regulates photosynthesis processes, while the pavement epidermis regulates growth responses to light perturbation. Heatmap diagram of DEGs (893 cluster-specific DEGs from Figure 7), grouped into blocks of expression patterns (left labels), comparing pavement epidermis and mesophyll cells (top label), across the continuing treatments (bottom labels). Table in right middle shows the weighted percentage overlap between clusters (Fig. 7) and blocks (this figure). Percentages are calculated by Jaccard method: Overlap / (cluster + block - overlap) x 100).

Across the dataset as a whole, we analysed the expression patterns in the epidermis and mesophyll for a curated set of light, signalling, clock and phyB-photobody associated genes **(Figure 9**, for relative expression data of all genes see **supplemental file 3)**. In the epidermis, PhyA and phyB are specifically upregulated at the end of the 4 hour dark treatment, while phyC,D and E expression is increased at +20 minutes red light. The PIF proteins have quite variable expression levels in the epidermis between treatments. However, in the mesophyll, photoreceptors and *PIF* expression levels are quite stable throughout. The scaffolding proteins *PCH1* and *TZP* have a differential regulation between epidermis and mesophyll **(Figure 9)**: *PCH1* is upregulated in the mesophyll at 12h dark, while *TZP* is more strongly regulated in the epidermis. *ELF3* is upregulated in the epidermis 4h Dark, while in the mesophyll it goes up at +20 minutes red. This shows that these factors, which have a known positive effect on phyB photobody stability, are regulated in a cell-specific manner. The circadian clock genes mostly follow expected day/night patterns, with subtle tissue-specific differences. Transcription factors that positively regulate photomorphogenesis are, like the PIFs, regulated more strongly in the epidermis **(Figure 9)**. The flowering genes *CO* and *FT* are mostly absent from the epidermis and upregulated in the mesophyll at 12h dark and +20 red respectively.

**Figure 9.**
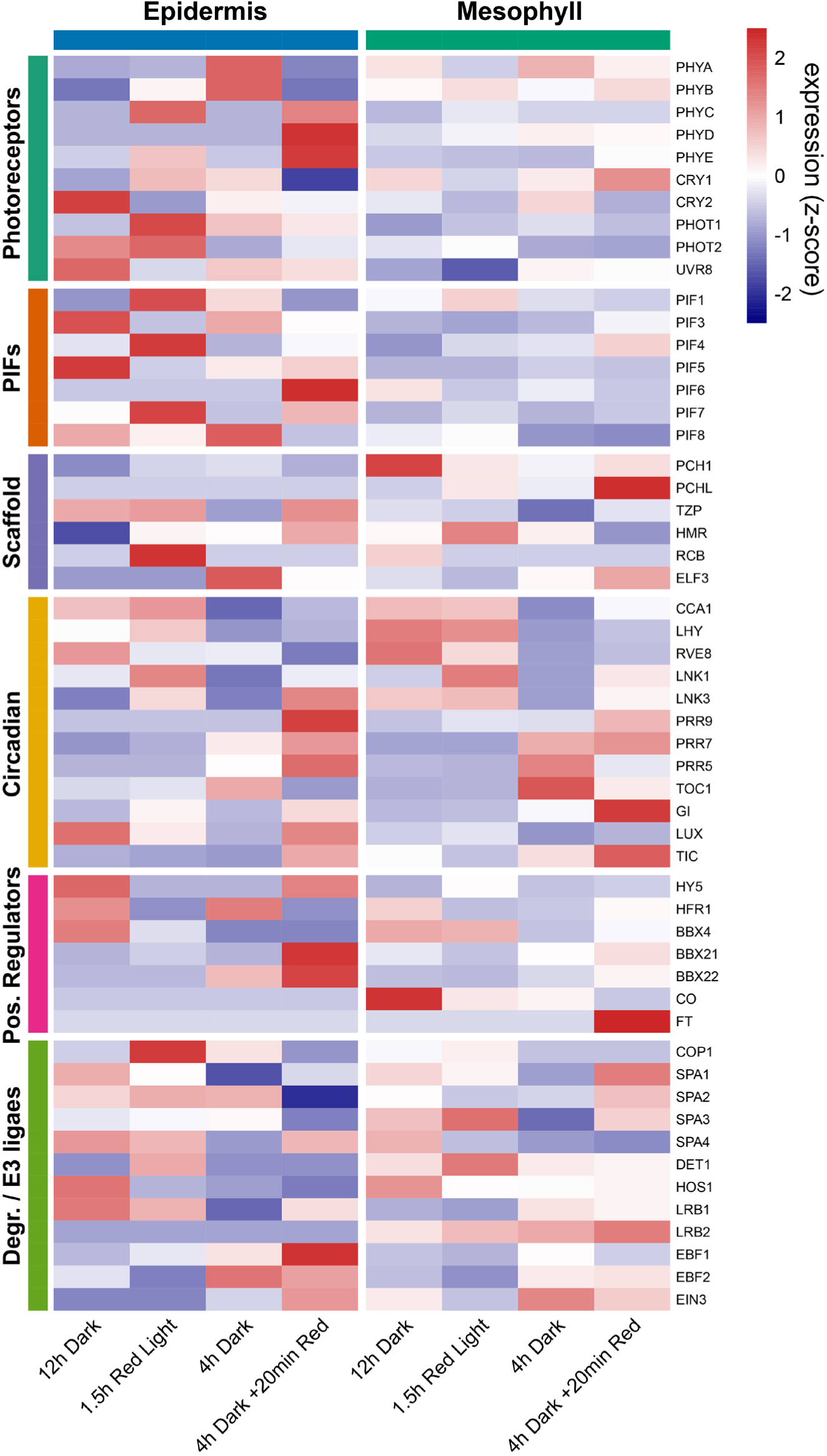
Tissue-and-light-specific expression patterns of photobody and light signalling genes show distinct epidermis and mesophyll roles in response to red light and darkness. Pseudobulk expression of a gene panel of across the four Col-0 treatments (12h Dark, 1.5h Red Light, 4h Dark, 4h Dark + 20 min Red Light) in epidermal and mesophyll nuclei. Per group, mean normalised expression was calculated across all nuclei and then z-scored per gene, reflecting relative expression between tissues and treatments. Genes are ordered by functional category: Photoreceptors, PIFs, phyB photobody promoting genes and scaffold components, core clock genes (Circadian), Positive regulators of light signalling (TFs), promoters of degradation (E3 ligases), and SPA kinases.

The light-regulatory E3 ligase complex consisting of *COP1* and *SPA* genes is differentially regulated across treatments and cell types, with COP1 being especially upregulated in the epidermis 1.5hr red **(Figure 9)**. *LRB1*, which controls PIF and phyB degradation (Ni et al. 2014), is upregulated in the epidermis at 12h dark and 1.5hr red, while *LRB2* is upregulated in the mesophyll at 4hr dark and +20 minutes red. Overall, the expression analysis patterns show differential regulation of expression clusters and light signaling genes between epidermis and mesophyll. The epidermis more strongly regulates light singling and growth responses to environmental perturbation, while the mesophyll regulates strongly regulates photosynthesis and circadian rhythms at the start of the day. Interestingly the largest block is Block 6, which is an epidermis-related set of growth regulation genes. The strongest overlap with photobody-related genes (cluster 4) can be found in the Block 1 and 2, which is regulated at 1.5h Red. At this timepoint, we also observed also a clear photobody response in all cell types, due to coarsening **(Figure 6B**, **8)**.

## Conclusions and discussion

Our experiments have shown that phyB photobodies exhibit coarsening behaviour, similarly to other bio-molecular condensates in plant and non-plant systems, but do not follow classical Ostwald ripening dynamics. Indeed, coarsening initially occurs relatively fast, and small photobodies disappear quickly. However, phyB photobodies have a limited size, and growth ceases prematurely, exhibiting signatures of arrested coarsening. One of the mechanisms that could lead to the arrest in coarsening is the nuclear organisation itself: there is evidence that some phyB photobodies associate with particular chromocenters (Du et al. 2024, 2026). The small photobodies we observe through super-resolution indicate a fine-grained network of potential nucleation sites for photobodies. Another possibility is that the arrest in coarsening reflects an interplay between condensate growth and non-equilibrium processes, such as light-dependent interconversion reactions or transcriptional regulation.

In Our FRAP experiments we found that certain cell types, such as the mesophyll have quite high FRAP recovery, up to 80%, while guard cells have a much lower, up to 30%. While our current FRAP setup has one important limitation, namely that bleaching of one photobody could lead to substantial bleaching of the overall pool of fluorescent proteins, we also find that different treatments and cell types exhibit different rates of recovery. In particular, darkness photobodies have slower recovery than photobodies in light.

In the single cell imaging experiment, we observed the fastest and strongest responses to light in the pavement epidermis cells, while the mesophyll palisade and spongy cells coarsened slower, and reacted slower to darkness. Interestingly their initial response to new light seemed to be of equal speed to the pavement cells. Intriguingly, the guard cells also responded less than the pavement cells, even though, they are in the same layer of cells. Overall, the mesophyll photobodies persisted longer through time, and were seemingly more stable, while at the same time showing the highest recovery rate in FRAP. These light and cell-type specific FRAP and response differences could be caused by several factors.

First, while it would make sense that cell-specific differences in photobody persistence are due to a different light penetration, this is likely not a major determinant here: Guard cells show higher photobody stability compared to the pavement cells, while receiving equal amounts of light. Furthermore, pavement cells are very transparent and have very few and small chloroplasts, making them quite transparent for light. As a result, palisade and spongy mesophyll beneath receive and absorb most of the light (Barton et al. 2016).

Second, light and cell-specific FRAP differences could be due to a change in phyB protein levels driven by light- and cell-specific expression differences, which has been shown before to be a determining factor (Chen et al. 2022; Kim et al. 2024). These changes in phyB protein levels are also quite clear when comparing our volume/intensity quantification between pavement cells to spongy and palisade cells in the single cell imaging series. Furthermore, transcriptionally, phyB is much higher expressed in the spongy mesophyll compared to other cell types **(Supplemental datafile 3),** and its transcriptional regulation is more stable in the mesophyll (**Figure 9)**. This observation is also interesting from a theoretical point of view since these changes in expression levels and/or the activity of phyB Pr vs Pfr might shift the system closer to or further away from the binodal line (in an equation of free energy in a phase separated system). Such a shift would change the timescale of molecular exchange and coarsening, potentially explaining the difference in FRAP recovery of fully bleached photobodies. Note that our FRAP experiments were performed with a native-expressing phyB line instead of with phyB overexpressing lines. This might explain why some of our FRAP data correspond to previously published literature, but that some does not. A higher FRAP recovery in lower red light, for instance, is consistent with previous work, where increased phyB expression correlates with a reduced FRAP rate and bigger photobodies (Kim et al. 2024). In contrast, we observed a higher FRAP rate in the larger, more highly expressed photobodies of the spongy mesophyll and palisade, compared to the smaller ones in the pavement cells. Thus, the size and intensity of photobodies does not always correlate with a raised or lowered FRAP rate, but there does seem to be a better correlation with phyB expression levels and persistence of photobodies in darkness, or far-red light. Overall, since we used a native-expressing line, we believe that our results should nicely reflect actual cell-type differences in growing plants, as compared to previous FRAP experiments.

A third important factor determining changes in photobody persistence and FRAP recovery rates are changes in interactions between phyB and other proteins or biomolecules in the photobody. Scaffolding proteins like PCH1, TZP and ELF3 are involved in creating a longer-persisting photobody, with evidence that phyB photobodies are less stable throughout darkness in knockout mutants of these genes (Enderle et al. 2017; Fang et al. 2022; Yang et al. 2025). During the night, interactions between phyB and these stabilizing factors can potentially strengthen oligomerization through interactions, which stabilize over time, reducing the number of binding and re-binding interaction events, thus ‘ageing’ the droplet, leading to persistence in the dark. When the light was applied after 12h Darkness, we observed a quick remodelling of phyB photobodies, as shown by the formation of new, smaller photobodies in all cell types, which could represent phyB being newly activated and establishing new interactions. Our single nucleus RNA expression analysis shows that each cell type has a different mix of stabilizing or scaffolding proteins being expressed, such as *PCH1, PCHL, TZP* and *ELF3*. *PCH1* is expressed strongly in the dark mesophyll, but not in the pavement cells, while *TZP* can be found more specifically in the pavement cells across conditions **(Figure 9)**. *TZP* was found to be part of the photobody-related Cluster 4, thus could contribute to conditional stability. However, with our current understanding of PCH1 and TZP, we cannot explain the differences in stability of pavement cell photobodies and mesophyll photobodies, since both have been shown to promote phyB photobody stability (Enderle et al. 2017; Huang et al. 2019; Fang et al. 2022).

Interestingly, phyB photochemistry, or the rate of active Pfr and inactive Pr does not seem to be correlated very well with detailed photobody dynamics. On the long run, photochemistry will prevail, as observed with a one-hour far-red treatment **(Figure 5)**. However, in our FRAP and single cell imaging results such a one-on-one correlation is lacking. In pavement cells, we found the highest FRAP recovery to be correlated to high red light, and thus active phyB, while this was the opposite in guard cells. Importantly, the second 20-minute light treatment led to a different response in photobodies, compared to the first red light treatment, even though this is a more than sufficient amount of time to elicit changes in photochemistry. Photochemistry can be altered very rapidly with far-red light, while 4 hours of darkness is enough to reduce most phyB to inactive Pr state (Rausenberger et al. 2010). That we still observed photobodies after 12h of darkness is also in line with existing literature (Buskirk et al. 2014; Enderle et al. 2017; Murcia et al. 2021), highlighting the disconnect between photochemistry and photobody formation, which is most likely being caused by the interaction with scaffolding proteins.

Complementing the imaging experiments, the single-nucleus transcriptomics experiment revealed that there are cell-specific roles of the pavement and mesophyll in responding to light, and absence of light. Both mesophyll and pavement epidermis cells respond by regulating the circadian clock at first light, and promoting amino acid and sugar catabolysis upon sudden darkness. Specific to the mesophyll is the regulation of photosynthesis at the start of the day, while the epidermis cells specifically regulate growth-promoting and restricting factors, especially in the second light application. Using *YHB* and *phyB* controls, in combination with nuclear-transcriptome-clustering analysis, we were able to identify a set of genes that are associated with the presence of photobodies. These include photosynthesis, circadian regulators, and mRNA binding, all processes previously associated with phyB-driven gene regulation, as is also evident from the overlap of the DEGs from this dataset with published papers (Sakuraba et al. 2014; Shikata et al. 2014; Huang et al. 2019; Kathare et al. 2022; Yan et al. 2022). Not surprisingly, there was a significant overlap between our differentially expressed genes and previous single nucleus leaf datasets (Tenorio Berrío et al. 2022; Xia et al. 2022; Qin et al. 2025).

Thus, phyB photobodies seem clearly associated with regulating transcription. We could identify a feedforward loop, where changes in phyB photobodies lead to changes in *TZP* expression, which can in turn stabilize phyB photobodies, leading to a ‘locking in’ of a particular light state, so that it can keep coarsening and persist in darkness, even when the photochemistry attempts to drive phyB into an inactive state. What is not clear form our analysis is whether the phyB photobody itself regulates transcription at the chromatin. phyB can elicit a Chromatin Immunoprecipitation signal (Jung et al. 2016), however, this could come from phyB either in our outside a photobody. PhyB-related transcription is mostly regulated via repression of PIF proteins, and PIFs can be observed in phyB photobodies when overexpressed. However, at native expression levels, PIFs are either not observed in photobodies, or hardly observable at all, due to their transient nature. Thus, only direct microscopical or biochemical observation will help answer the question of direct involvement of the photobody in transcription. The other mechanism through which photobody memory affects transcription and light signaling is by the condensation and inhibition of signaling molecules present in the photobody itself. Thus, by continued inhibition of specific regulatory factors, or releasing them at a particularly delayed time, phyB molecular memory can potentially smooth-out signaling events due to changing light conditions, which might otherwise be too costly to constantly reprogram. However, when the plant needs to respond to an important stimulus, such as dawn, it will do so. Thus, the circadian clock can regulate factors affecting phyB photobodies, while light detection through phyB can set the clock. If a perturbation then occurs throughout the day, phyB photobody memory allows the plant to partially ignore this signal, or induce growth regulation to escape the stress. Only a consistent signal, such as a longer or intense amount of far-red light, will then elicit a photobody response.

To conclude, we have observed that phyB photobodies display molecular memory that can be influenced by different light treatments and that differs between cell types. Conditional-specific phyB responses have been characterized before in the context of increased temperature, where phyB together with PCH1 can store night-time information (Murcia et al. 2021). In this study, we observe that phyB photobodies also display a conditional-specific persistence, and arrested coarsening behaviour. Transcriptional, cell-specific and treatment specific responses are different after repeated stimuli and between cell types. Thus, cell-biologically, and transcriptionally, phyB photobodies display memory function. A potential mechanism for this phenomenon is the cell-and light specific expression of photobody-stabilizing factors such as TZP and PCH1, however, their cell-specific roles, or differences in molecular function are less clear. More broadly, these observations suggest that phyB photobody memory can add a crucial extra layer to light regulation, which allows the plant to respond consistently to changing environments. Furthermore, photobody dynamics might allow plants to integrate multiple environmental inputs such as light and temperature rather than reflecting just the current light condition. Experiments varying light and temperature independently and simultaneously could test such processing of multiple signals and might help elucidate whether photobody dynamics contributes to robust information processing.

## Materials and Methods

### Genetic material, plant growth and growth conditions

Lines used in this study were Col-0, *phyB-9* (Reed et al. 1993), *YHB* (Su and Lagarias 2007), and *phyB-9 pphyB:phyB-YFP* (Viczián et al. 2020). Seeds were surface sterilized by sequential washing in 70% ethanol. Approximately 1.5 ml of ethanol was added to the seed bag to fully soak the seeds, the bag was inverted, and an additional 1.5 ml of ethanol was added. After drying in a fume hood, the process was repeated once more before seeds were sown onto ½ MS plates. Plates with seeds were stratified for 3 days. All seedlings were grown initially in white light conditions at 21°C for five days, after which the respective light treatments were applied **(Figure S14)**. White light was applied through Valoya white LEDs with a PPFD of 145.4 µmol m⁻² s⁻¹ and a red light strength (600-700 nm) of 60 µmol m⁻² s⁻¹. Red LED light of 660 nm was applied at 60 µmol m⁻² s⁻¹ for the high-red light treatment, and at 15 µmol m⁻² s⁻¹ for low-red. Experiments were carried out in a closed chamber with darkness values of (PPFD = 0.0019 μmol m⁻² s⁻¹; PFD-UV = 0.0005; PFD-G = 0.0006; PFD-B = 0.0005; PFD-FR = 0.0009; PFD-R = 0.0008). To avoid influences of the circadian clock, all experiments were carried out at fixed times during the day. Seed sowing and stratification at 16:00 on the day of setup. Plants were put in growth chambers at 10:00. Overnight treatments were initiated at 20:00 and maintained until 08:00 the following morning, after which the respective light treatments were applied. All harvesting and imaging procedures were conducted after 08:00 to maintain temporal consistency across experiments. Light spectra were measured with a UPRtek PGN200 spectrophotometer.

### Confocal sample preparation and fixation

Live imaging samples were prepared by cutting a cotyledon of a seedlings, and placing it on a small coverslip-bottomed cell chamber, with a droplet of water and a 1.5% agarose block placed on top. For differential light conditions during imaging, flexible amazon USB reading lights were modified with different LEDs (white, 660, 730 nm) on top, and were placed in a generic USB hub with on/off switches. The LEDs were aimed at the sample during confocal imaging. Light intensities were measured with a mock setup using the spectrophotometer. The condenser of the transmitted light path of the microscope was shielded with a green filter by Lee.

The samples for cell-specific photobody analysis were fixed in 4% PFA in their respective light treatment for 10 minutes, followed by vacuum application for 20 minutes in darkness, followed by 1 hour of room temp fixation at normal pressure. Samples were washed twice with 1x PBS buffer, after which clearsee alpha was applied for two weeks (Kurihara et al. 2021). One day prior to mounting, seedlings were stained with a solution of 0.01% calcofluor white / fluorescent brightener 28, to stain cell walls, overnight, followed by a 30 minute clearsee wash (Ursache et al. 2018). Seedlings were mounted on slides with minimal fresh clearsee with 32 mm cover slips and left overnight to seal.

### Confocal microscopy

Confocal imaging was performed on a Leica Stellaris SP8, and a Zeiss LSM 880 Airyscan. Live imaging parameters on the SP8 Stellaris: Objective, HC PL APO CS2 63×/1.20 WATER; dimensions and pixel size 512×512×20, 0.212 µm; laser, 515 nm 4% power; detector, hyDX (520-560) gain 350, speed 400Hz bidirectional. Images were taken at 5-minute intervals to minimize laser exposure.

FRAP live imaging parameters: SP8 Stellaris: Objective, HC PL APO CS2 63×/1.20 WATER, specialized LEICA FRAP lens was used; dimensions and pixel size 512×512×13, 0.048 µm; laser, 515 nm 6% power; detector, hyDS (519-579) gain 350.

Cell-specific photobody imaging setting on Zeiss LSM880 Airyscan: Objective, Plan-Apochromat 63×/1.40 Oil DIC M27; dimensions and pixel size 956×956×50 0.045 µm; Laser 514 15% equivalent; Detector Airyscan detector Fast Mode, 516-579 nm.

### Image processing, segmentation, and data analysis

Images were processed using ICY (De Chaumont et al. 2012) for figure preparation. phyB photobodies were 3D segmented using Biom3D https://biom3d.readthedocs.io/ (Mougeot et al. 2026). A deep learning Biom3D segmentation model was trained using 100 individual confocal image slices annoted using Napari https://napari.org/. A custom python script was written to process time series confocal images and to extract data from Napari using masks generated by Biom3D. For more details see **Supplemental methods**.

FRAP images were segmented using HK means of ICY for Figure 3. For Figure 4 we developed a FRAP 3D tracking method called dynaFRAP https://github.com/andreweuw/DynaFRAP. dynaFRAP uses the aforementioned biom3D method to segment photobodies, to then use this segmentation to follow and determine FRAP-ed regions through time and XYZ-space. dynaFRAP contains a built-in post-processing analysis and plotting mode. For more details see **Supplemental methods**.

Unless otherwise stated, Imaging data was analysed using R and Rstudio.

### Nuclei isolation protocol

Seedlings were placed in rows on ½ MS agar plates and grown as indicated. Plates were flash frozen whole in liquid nitrogen, and placed in a -80°C freezer. Then, cotyledons were scraped off into fresh liquid nitrogen using a scalpel. 2 grams of frozen cotyledons were ground for approximately 6 minutes in a potter homogenizer together with 2ml of ice-cold nuclei isolation buffer (For buffer compositions, see below). Homogenized cotyledons were filtered sequentially through 40 and 20 µm pre-wetted Pluri Select Cell strainers (Cat. # 43-50020-03 and 43-50040-50) and and centrifuged at 100 g for 1 minute at 4°C. Flow through was then spun down at 1000 g for 10 minutes at 4°C and resuspended in 500 μL Nuclei Isolation Buffer. This was carefully layered onto a Percoll–sucrose density gradient (1:1:1 mix of 2.3 M sucrose, 30% Percoll, sample respectively). Percoll gradient was then centrifuged at 1,000 g for 30 min at 4°C. Nuclei were then collected from the green interphase using cut 200 μl tips (Around 500 μL). The nuclei fraction was then transferred to 2 mL tubes and diluted with 1.5 mL Nuclei Resuspension Buffer, and subsequently spun down at 1000 g for 10 minutes at 4°C. The pellet was then washed with 2 mL Nuclei Resuspension Buffer and spun down again, resuspended in 500 μL Nuclei Resuspension Buffer and transfer to precooled 15 mL tubes. 2 mL methanol + 150 μL BS3 crosslinker (Thermo Fisher cat. no. 21580) was added drop by drop on ice and incubated for 15 minutes on ice. Nuclei were then resuspended in 4 mL of Resuspension Buffer. The mixture was then centrifuged at 800 g for 7 minutes at 4°C, then pellet was resuspended in 1mL Resuspension Buffer. The nuclei were counted and QC-ed using a LUNA-FX7™ Automated Cell Counter (Logos Biosystems, South Korea).

*Nuclei isolation buffers.* All solutions were prepared in nuclease-free water. A 10× hypotonic PBS stock (5.45 g Na₂HPO₄, 3.1 g NaH₂PO₄·H₂O, 1.2 g KH₂PO₄, 1 g KCl, 3 g NaCl per 500 mL) was stored at room temperature for up to 6 months. Base hypotonic lysis buffer consisted of 1× hypotonic PBS supplemented with 3 mM MgCl₂ and was stored at 4 °C. Working Nuclei Isolation Buffer was prepared fresh and kept on ice, and consisted of base lysis buffer containing 0.8 mg/mL BSA, 0.05% (vol/vol) Igepal CA-630 and 1% (vol/vol) DEPC. Nuclei Resuspension Buffer (0.3 M SPBSTM) contained 0.33 M sucrose, 1× dPBS, 0.1% (vol/vol) Triton X-100 and 3 mM MgCl₂; it was sterile-filtered and stored at 4 °C for up to 3 months. BS3 crosslinker was reconstituted in 1 mL of 0.3 M SPBSTM and diluted into a further 2 mL of the same buffer (3 mL final), aliquoted at 150–300 µL and stored at −80 °C for up to 3 months; aliquots were thawed once and not refrozen.

### Single Cell Combinatorial Indexing RNA Sequencing (sci-RNA-seq3)

The sci-Seq protocol and library prep was then performed according to the method of (Martin et al. 2023), with the following modification: Unless stated otherwise, nuclei were pelleted at 700 g for 3 min (rather than 500 g). Thawed aliquots were washed in 200 µl 0.3 M SPBSTM and sonicated routinely (12 s, low, Bioruptor; ≤300 µl per tube) rather than only when visibly clumped. Nuclei were counted after SYTOX staining (7 µl SPBSTM, 1 µl SYTOX, 2 µl nuclei) instead of Yoyo-1. Primer and reagent plates were spun at 1,000 g for 1 min after removal from 4 °C to avoid microdroplet carry-over. Nuclei were loaded at 25,000 per well (200,000 per column, 2.4 × 10^6^ per plate) versus 2 × 10^6^ per plate; the nuclei/dNTP table was extended to a half-column (100,000 nuclei, 21.25 µl, 2.4 µl 10 mM dNTPs). Reverse transcription was performed with Maxima H Minus reverse transcriptase (0.5 µl enzyme, 2 µl 5× buffer, 0.5 µl water per well) in place of SuperScript IV, and using a stepped ramp — 4, 10, 20, 30, 40 and 50 °C for 2 min each, followed by 55 °C for 15 min (lid 65 °C) — instead of a single 55 °C, 10 min incubation. Wells were pooled by row with a 12-channel pipette set to 15 µl. In the ligation, after the final washes, nuclei were resuspended in 200 µl (rather than 1 ml) SPBSTM, and the post-ligation count was additionally used to estimate the doublet rate. For second strand synthesis, nuclei were distributed at 5 µl per well of a standard 96-well plate into second-strand synthesis mix prepared as a single master mix (488.75 µl water, 57.5 µl 10× buffer, 28.75 µl enzyme per plate). Incubation was 2.5 h at 16 °C or overnight at 4 °C. Prior to library tagmentation, a tagmentation test was performed, which is not part of the published protocol. A twofold serial dilution series of N7-loaded Tn5 in TD buffer (1×, 2×, 4×, 8×) was applied in duplicate to test wells (5 µl per well, 55 °C for 5 min), followed by transposase removal (0.4 µl 1% SDS, 0.4 µl BSA, 1.8 µl water; 55 °C for 15 min), quenching with 2 µl 10% Tween-20, and 16 cycles of PCR. Products were purified with 0.8× SPRI beads, eluted in 10 µl EB and quantified by Qubit dsDNA HS assay and D1000 HS TapeStation. The optimal Tn5 concentration was selected from this titration and used for the library plate, in place of the fixed 0.125 µl N7-loaded Tn5 per well specified in the published protocol. Tagmentation, transposase removal and quenching were as published. Third-round indexing was applied as a dual-index combination — 2 µl of indexed P5 primer per well plus 2 µl of row-specific indexed P7 primer carried in the master mix (20 µl 2× NEBNext, 2.4 µl water; 24.4 µl per well) — rather than 96 uniquely indexed P7 primers with a single P5 primer included in the master mix at 0.2 µl. Wells were pooled and a 300 µl aliquot taken forward, with the remainder stored at −20 °C. The library was purified with 0.8× SPRI beads, washed twice with 80% ethanol (rather than 70%) and eluted in 30 µl EB. The PAGE check of individual wells, the preparative 1% agarose gel and the 250–600 bp gel extraction described in the published protocol were omitted; final libraries were instead quantified by Qubit dsDNA BR assay and sized on a D1000 BR TapeStation, and diluted to 4 nM for sequencing. The libraries were sequenced with the NextSeq2000 P2 kit.

### sci-SEQ data analysis

Reads were demultiplexed, trimmed, aligned to the Arabidopsis thaliana TAIR10 genome (UTR-extended annotation) and UMI-counted per gene over exons and introns with the sci-rocket workflow (https://github.com/lauren-saunders-lab/sci-rocket;(Panten et al. 2024)) using STARsolo (Kaminow et al. 2021), yielding 3,621 nuclei across the six samples. Downstream analysis was performed in R with Seurat v5 (Hao et al. 2024). Genes detected in fewer than three nuclei and nuclei with fewer than 200 genes were removed (2,241 nuclei × 21,102 genes), and nuclei with >2,500 genes, >10,000 UMIs, >10% chloroplast (ATCG) or >5% mitochondrial (ATMG) reads were excluded as multiplets or ambient-RNA-contaminated nuclei (1,884 nuclei; **Figure S7**). Counts were log-normalised (scale factor 10,000), 3,000 variable genes were selected (vst; organellar genes excluded) and, after scaling, the first 15 principal components were used for shared-nearest-neighbour graph construction (k = 20), Louvain clustering and UMAP; no batch correction was applied because all samples were intermixed. Two small clusters (101 nuclei) with low UMI depth and no coherent markers, derived only from the two 4h-Dark treated samples, were removed as low-quality nuclei and the analysis was repeated on the remaining 1,783 nuclei, with the final resolution (0.3, eight clusters) chosen with clustree (Zappia and Oshlack 2018). Cluster-specific DEGs were identified with FindAllMarkers (Wilcoxon rank-sum test, min.pct 0.25, |log2FC| ≥ 0.5, Bonferroni-adjusted p < 0.05), and cell types were assigned per cluster from module scores (AddModuleScore). Treatment-responsive genes were obtained from all six pairwise FindMarkers comparisons between the four Col-0 conditions (min.pct 0.1, |log2FC| ≥ 0.25, Bonferroni-adjusted p < 0.05; 739 unique DEGs), and their mean normalised expression per tissue (epidermis, mesophyll) and condition was z-scored and hierarchically clustered (Euclidean distance, Ward’s method; pheatmap) into expression blocks **(Figure 8)**. GO (Biological Process) enrichment of clusters and blocks was performed with clusterProfiler (Wu et al. 2021) and org.At.tair.db, using all 21,102 detected genes as background and Benjamini–Hochberg-adjusted p < 0.05; overlaps with published gene sets **(Figure S12)** were tested by Fisher’s exact test against the same background with Benjamini–Hochberg correction. Gene identities were verified with org.At.tair.db (TAIR10; R [x.y.z], Seurat [x], clusterProfiler [x]). As each condition was represented by a single sample, nuclei served as replicates and P values should be regarded as descriptive.

## Supplemental methods

Description of Biom3D pre- and post-processing code and dynaFRAP code development.

## Supplemental files

Supplemental File 1, all DEGs specifying nuclei clusters (Accompanying Figure 7)

Supplemental File 2, Genes and GO terms of expression pattern blocks (Accompanying Figure 8)

Supplemental File 3, Pseudobulk expression values of all epidermis and mesophyll genes.

## Data availability

Raw sequencing data of the sci-seq described in this study is deposited at the ArrayExpress of EMBL-EBI under project accession E-MTAB-17871.

## Declaration of generative AI and AI-assisted technologies in the manuscript preparation process

During the preparation of this work, the authors used Claude AI in order to assist with the python coding for differential gene expression analysis of the sci-seq data and to compare the DEG dataset of supplementary file 1 against known literature to generate the data for supplementary Figure 13. The authors reviewed and edited the content as needed and take full responsibility for the content of the published article.

## Author Contributions

F.I., L.W., L.M.S., I.G. and K.v.G. designed the study. F.I., F.S., S.H., J.S., M.C. and O.H. performed experiments. F.I., L.W., F.S., S.H., O.H. J.S., I.Z. and K.v.G. analysed data, and I.Z. and O.H. wrote new program code to analyse data. F.I., L.W., L.M.S., I.G. and K.v.G. wrote the manuscript.

**The authors declare no competing interests.**

