## Supplemental Figures for "phyB photobodies display molecular memory, regulating light signaling transcriptional states through phase separation"

### Supplemental data:

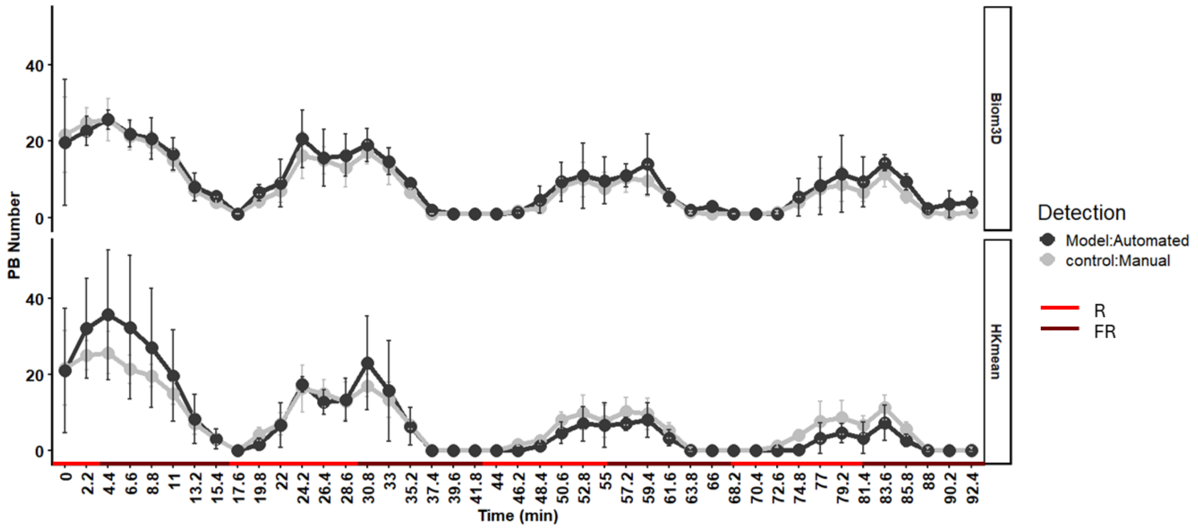


**Supplemental Figure 1: Biom3D is equally accurate to hand counting.** Comparison of average photobody number per nucleus quantification between a trained Biom3D model, and a Clustering and thresholding algorithm 'HK-Means' from ICY bioimage analysis vs hand counting of photobodies. Data gathered from a confocal live imaging experiment of *pphyB:phyB-YFP* seedlings exposed to alternating red and far-red light using LEDs. Upper panel = biom3D vs hand counting, lower panel = HK means vs hand counting. error bars = SEM, n = 10.

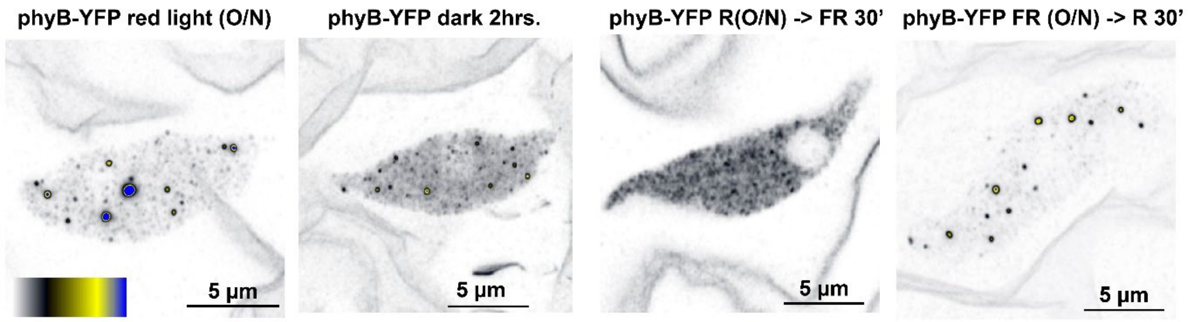


**Supplemental Figure 2. The distribution of phyB photobodies at super-resolution.** (A) Pavement cell nuclei of clearsee-fixed seedlings treated with red light (660 nm, 30 µE), far-red light (730 nm, 30 µE), and darkness, either overnight (O/N), 30 minutes, or 2 hrs, imaged with Zeiss Airyscan.

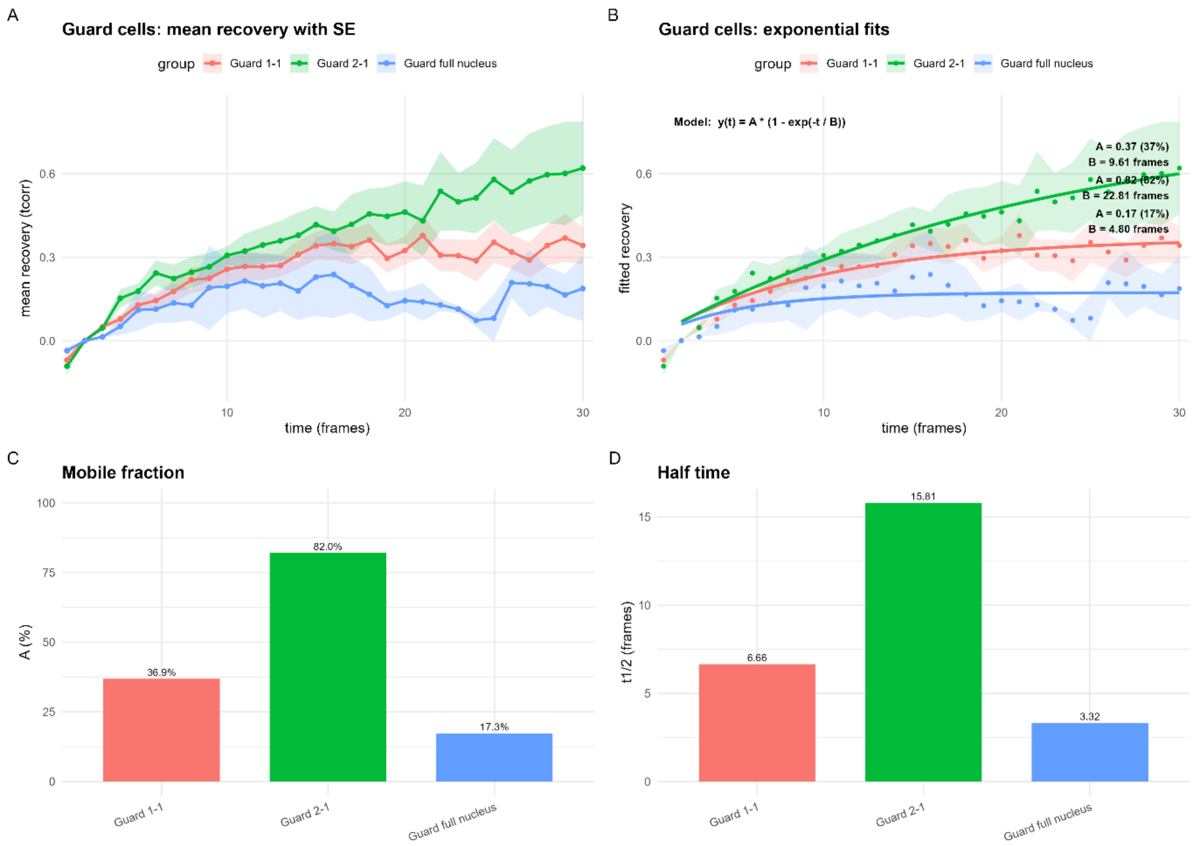


**Supplemental Figure 3. FRAP recovery of photobodies in guard cells with different bleaching strategies.** (A) Mean recovery curves with standard error (shaded areas) for three bleaching strategies: Guard 1–1 (Guard cell with a single photobody, which was bleached, red line), Guard 2–1 (Guard cell with two or more photobodies, with one photobody bleached, green line), and Guard full nucleus (entire nucleoplasm including photobodies bleached, blue). (B) Exponential fits of the recovery curves for each strategy, with fitted parameters shown (mobile fraction A and halftime B). (C) Bar chart showing the fitted mobile fraction (%) for each strategy. (D) Bar chart showing the recovery halftime (t½, in frames) for each strategy.

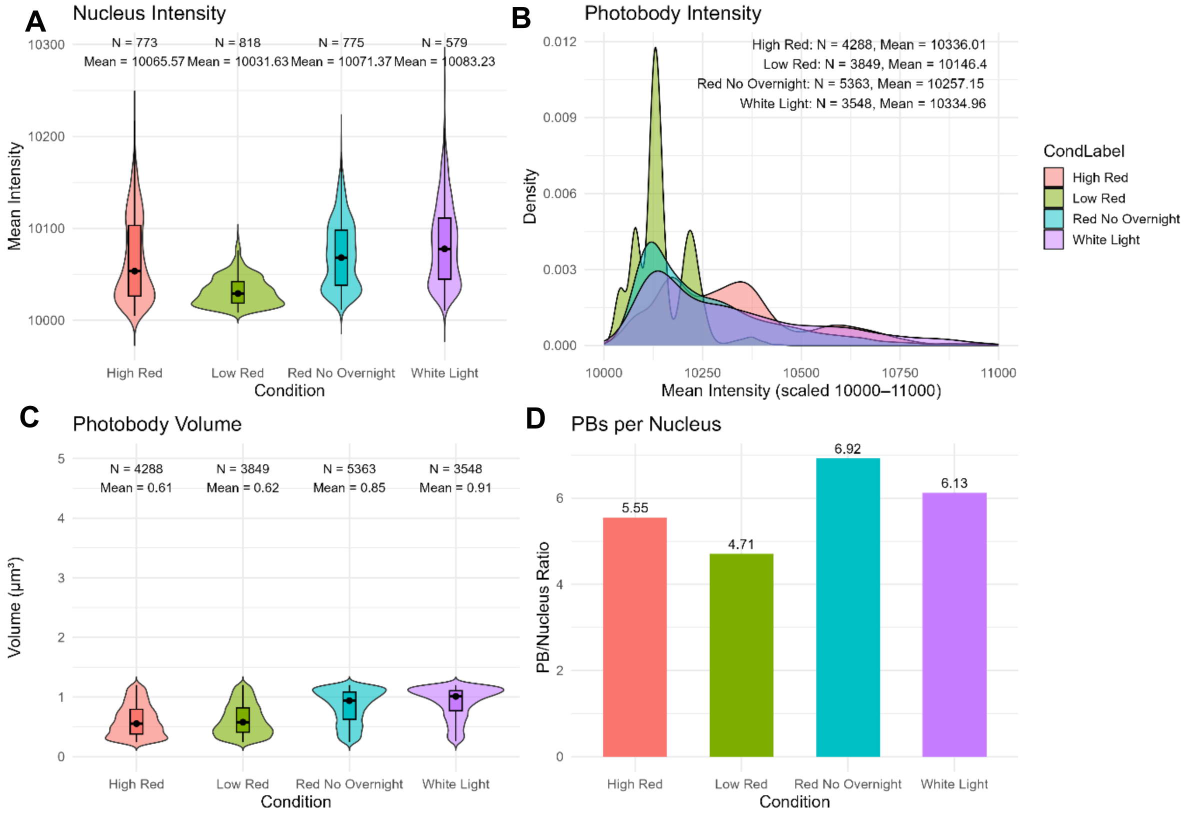


**Supplemental Figure 4: Light specific photobody measurements in *Arabidopsis* cotyledons.** Quantified parameters of cotyledon images of fixed seedlings, analysed with Biom3D, in high red (660 nm, 60 µE.), low red (15 µE), Hihg red with no overnight darkness (red no overnight), and White Light (PPFD 140 60 µE). (A) Average intensity of nuclei (arbitrary units, 10000 is equal to 0 due to airyscan processing algorithm. (B) Intensity frequency distribution of photobodies. (C) Average photobody volume. (D) Photobodies per nucleus ratio.

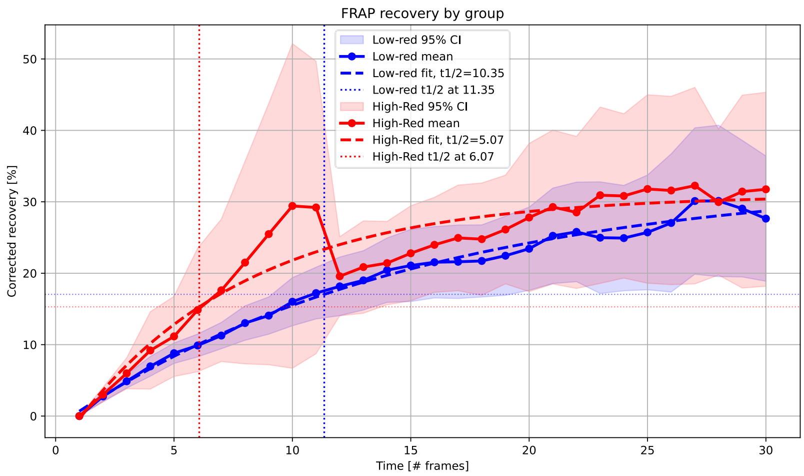


**Supplemental Figure 5. Guard cell FRAP in low red and high red light shows a difference in half time recovery, but not mobile fraction.** (A) FRAP experiment with guard cells in low (15 µmol/$m^2$/s) or high (60 µmol/$m^2$/s) red light (660 nm).

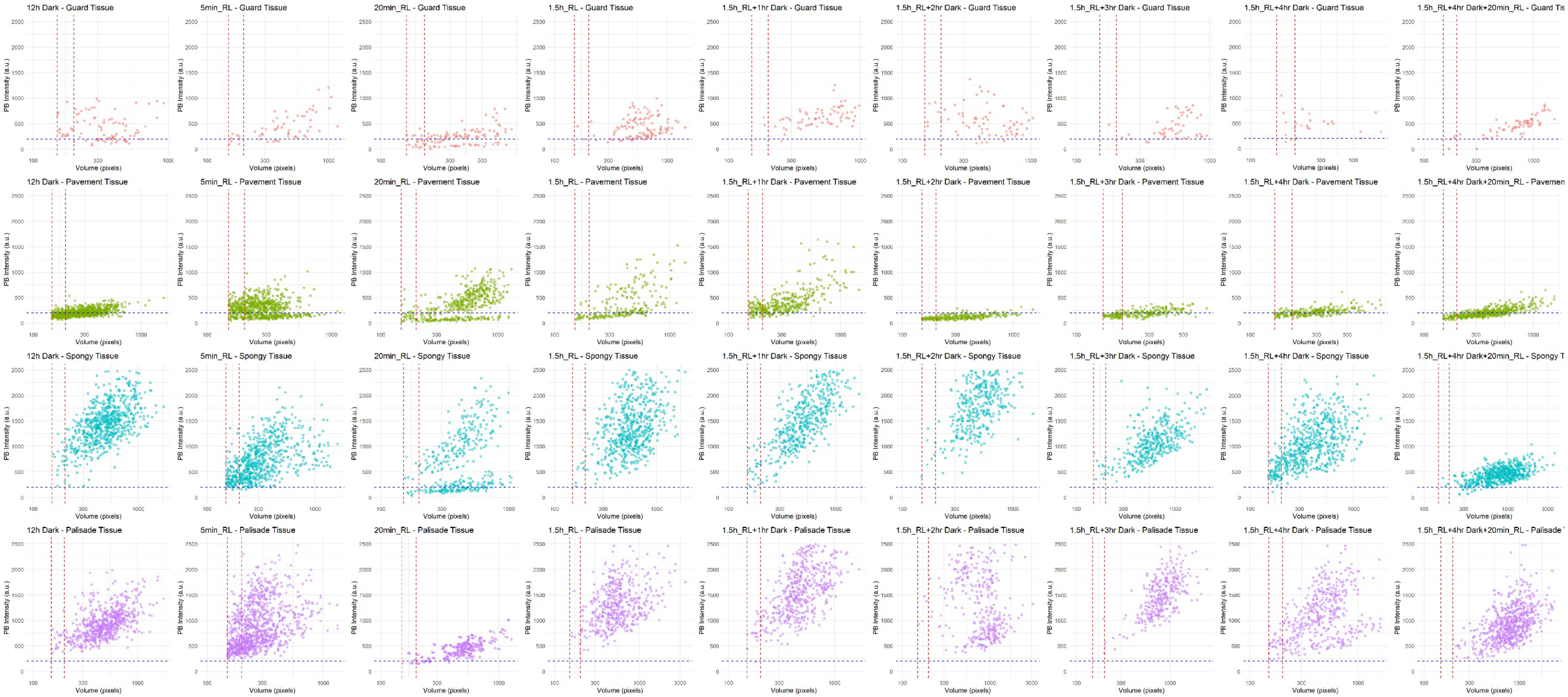

12h Dark - Guard Tissue
5min_RL - Guard Tissue
20min_RL - Guard Tissue
1.5h_RL - Guard Tissue
1.5h_RL+1hr Dark - Guard Tissue
1.5h_RL+2hr Dark - Guard Tissue
1.5h_RL+3hr Dark - Guard Tissue
1.5h_RL+4hr Dark - Guard Tissue
1.5h_RL+4hr Dark+20min_RL - Guard Tis
12h Dark - Pavement Tissue
5min_RL - Pavement Tissue
20min_RL - Pavement Tissue
1.5h_RL - Pavement Tissue
1.5h_RL+1hr Dark - Pavement Tissue
1.5h_RL+2hr Dark - Pavement Tissue
1.5h_RL+3hr Dark - Pavement Tissue
1.5h_RL+4hr Dark - Pavement Tissue
1.5h_RL+4hr Dark+20min_RL - Pavemen
12h Dark - Spongy Tissue
5min_RL - Spongy Tissue
20min_RL - Spongy Tissue
1.5h_RL - Spongy Tissue
1.5h_RL+1hr Dark - Spongy Tissue
1.5h_RL+2hr Dark - Spongy Tissue
1.5h_RL+3hr Dark - Spongy Tissue
1.5h_RL+4hr Dark - Spongy Tissue
1.5h_RL+4hr Dark+20min_RL - Spongy T
12h Dark - Palisade Tissue
5min_RL - Palisade Tissue
20min_RL - Palisade Tissue
1.5h_RL - Palisade Tissue
1.5h_RL+1hr Dark - Palisade Tissue
1.5h_RL+2hr Dark - Palisade Tissue
1.5h_RL+3hr Dark - Palisade Tissue
1.5h_RL+4hr Dark - Palisade Tissue
1.5h_RL+4hr Dark+20min_RL - Palisade
PB Intensity (a.u.)
Volume (pixels)

**Supplemental Figure 6: Individual scatterplots reveal tissue-specific responses to light and preservation of previous light states (Accompanying Figure 6).**

Quantified data of photobody intensity and volume taken from confocal microscopy with Airyscan super resolution module images of individual photobodies across four tissue types (Figure 6, Guard, Pavement, Spongy, and Palisade), under selected light and dark treatments (12 h Dark, 5 min Red Light, 20 min Red Light, 1.5 h Red Light, 1.5 h Red Light + 1 h Dark, 1.5 h Red Light + 4 h Dark, and 1.5 h Red Light + 4 h Dark + 20 min Red Light). Each point represents a single annotated photobody. The dashed red vertical line indicates the applied threshold for photobody volume (10200 au.), while the dashed blue horizontal line indicates the threshold for photobody intensity. These cut-offs were used to exclude immature or unstable condensates and segmentation artefacts.

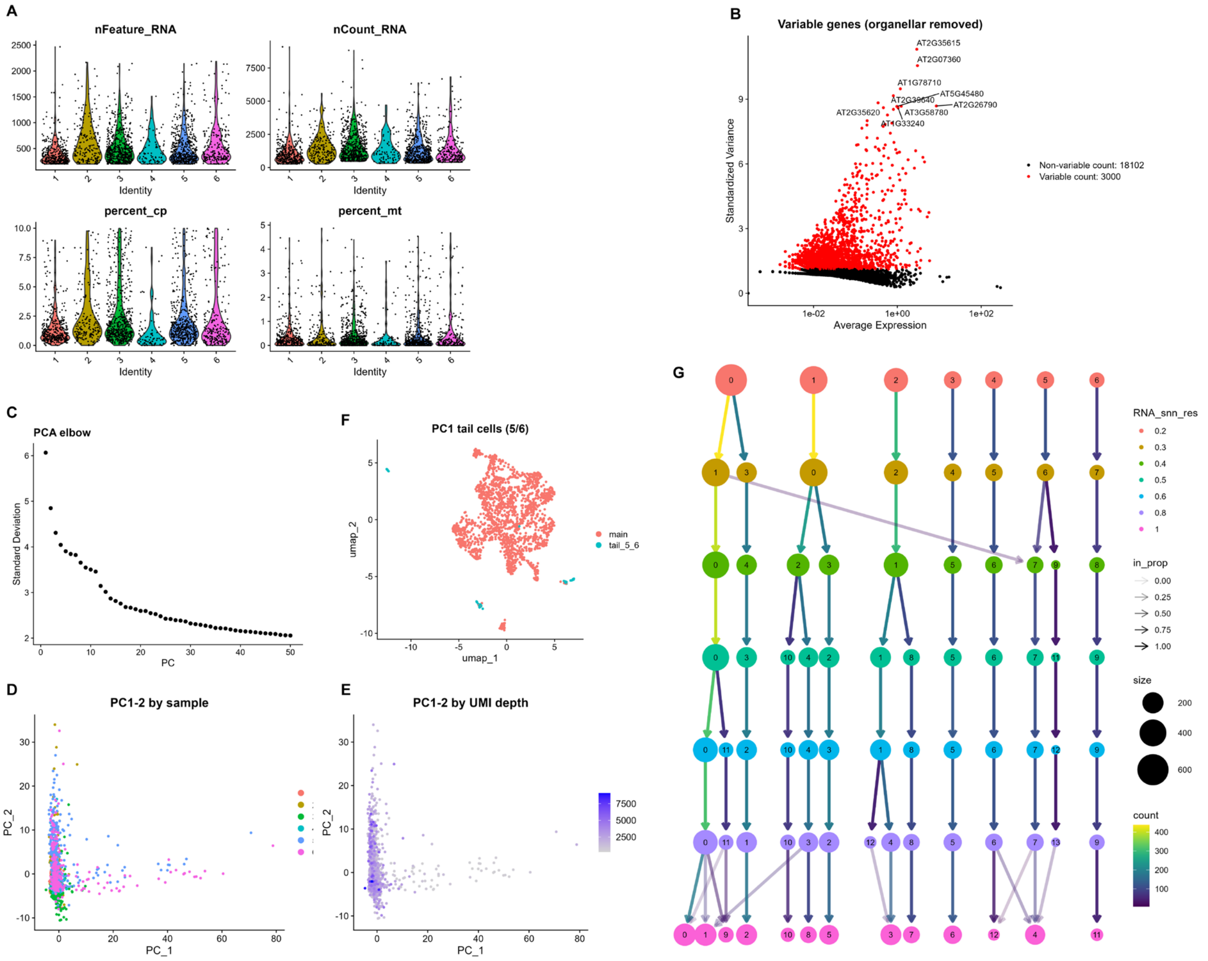


**Supplemental Figure 7. Quality control of snRNAseq dataset and Cluster analysis.**
(A) Distribution of the number of detected genes, the number of UMIs, and the proportion of chloroplast- (percent_cp) and mitochondrion-encoded (percent_mt) transcripts per nucleus for each of the six samples after filtering. Nuclei were retained with 200–2,500 detected genes, ≤10,000 UMIs, ≤10% chloroplast and ≤5% mitochondrial reads; the upper gene and UMI thresholds correspond to approximately the 99th percentile of each distribution and remove likely multiplets, while the chloroplast threshold removes nuclei carrying ambient plastid RNA. Of the 3,621 nuclei recovered by the sequencing pipeline, 2,241 passed the initial gene and cell filters and 1,884 passed quality control. (B) Mean–variance relationship of all detected genes; the 3,000 genes with the highest standardised variance (red) were selected for dimensionality reduction, after exclusion of chloroplast- and mitochondrion-encoded genes, which otherwise dominate the leading principal components. The ten most variable genes are labelled. (C) Standard deviation explained by the first 50 principal components; the first 15 components were retained for all subsequent analyses. (D, E) Nuclei projected onto the first two principal components and coloured by sample of origin (D) or by sequencing depth (E), showing that samples intermix and that depth does not drive the leading components; no batch correction or covariate regression was therefore applied. (F) UMAP embedding highlighting a group of 101 nuclei forming a tail along PC1, originating predominantly from the two 4 h dark-treated samples. These nuclei had approximately half the median sequencing depth of the remaining nuclei and no coherent marker gene expression, and were removed as low quality, leaving 1,783 nuclei. (G) Clustering tree across clustering resolutions 0.2–1.0. Node colour indicates resolution, node size the number of nuclei, and arrow transparency the proportion of nuclei moving between clusters. Resolution 0.3, yielding eight clusters, was selected as the highest resolution at which clusters remain stable before fragmenting.

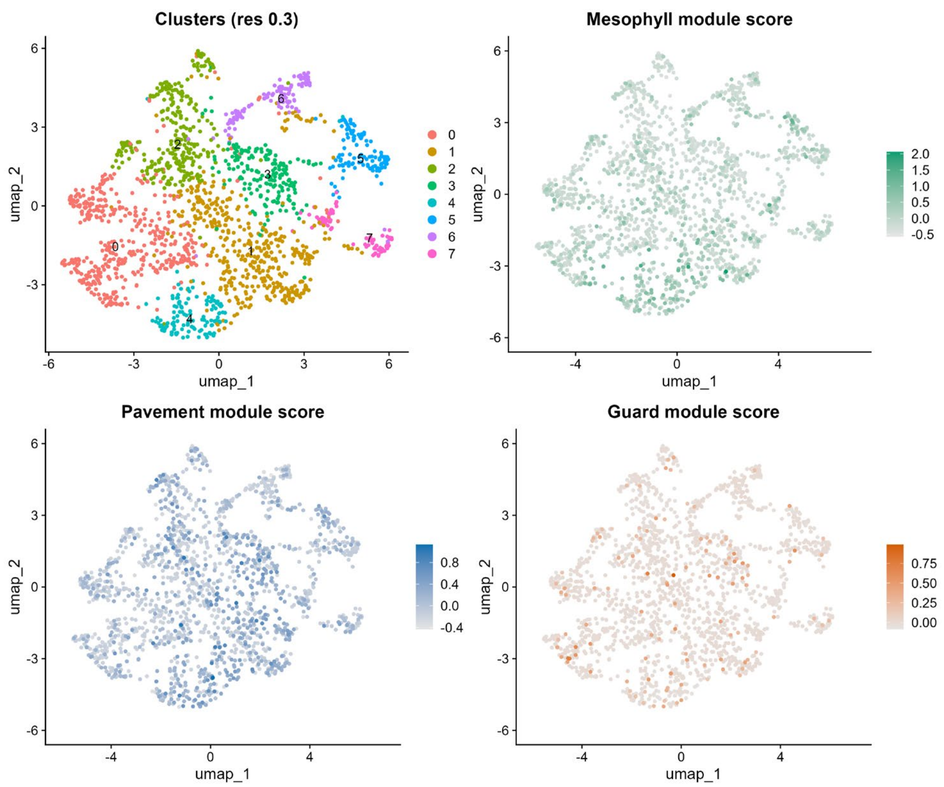


**Supplemental Figure 8. Cell type specification across the snRNAseq dataset**

UMAP embedding of all 1,783 nuclei coloured by cluster identity at resolution 0.3 (upper left), and by module scores calculated with cell-type marker panels from the *Arabidopsis* cotyledon single-cell atlas (Lee et al. 2025) for mesophyll, pavement (epidermis) and guard cells. Module scores were computed with AddModuleScore against a background of 100 randomly selected control genes matched for expression level. The mesophyll signature is broadly distributed across the embedding, consistent with most clusters representing mesophyll-derived cell states rather than distinct cell types. The guard-cell signature is scattered across individual nuclei without marking any cluster, indicating that guard cells are present but too rare to cluster separately.

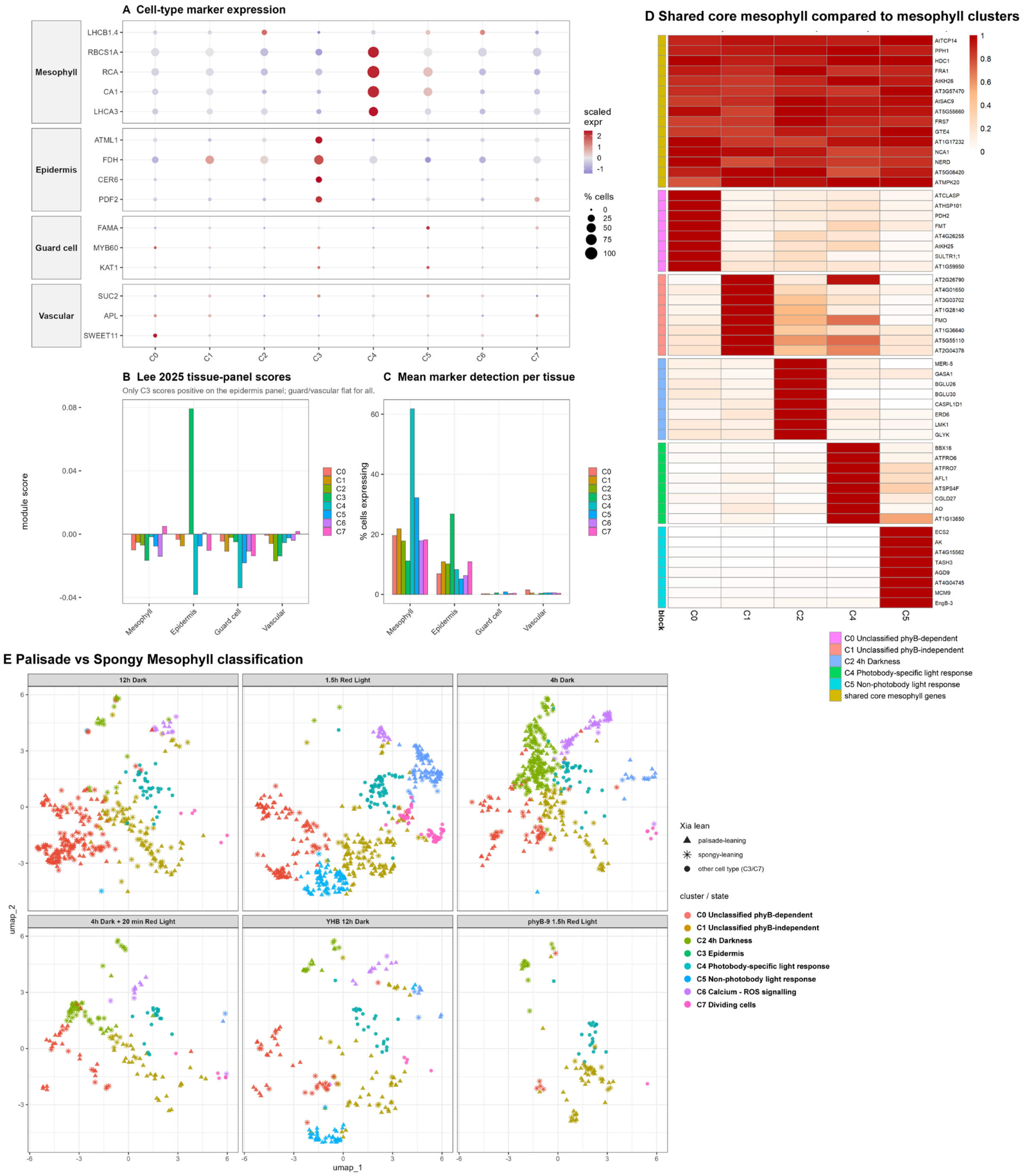


**Supplemental Figure 9. Mesophyll and epidermis can be confidently separated. Vasculature and guard cells are low in abundance. Palisade and Spongy Mesophyll have overlapping signatures.** (A) Expression of canonical cell-type marker genes across the eight clusters. Dot size indicates the percentage of nuclei in which a gene is detected and colour the mean expression scaled across clusters. Mesophyll markers (*LHCB1.4*, *RBCS1A*, *RCA*, *CA1*, *LHCA3*) are broadly expressed and highest in Cluster 4, epidermal markers (*ATML1*, *FDH*, *CER6*, *PDF2*) are restricted to Cluster 3, whereas guard-cell (*FAMA*, *MYB60*, *KAT1*) and vascular (*SUC2*, *APL*, *SWEET11*) markers are detected only in scattered individual nuclei. (B) Module scores of the tissue-specific marker panels of Lee et al. (2025) per cluster. Only Cluster 3 scores positively on the epidermis panel, while the guard-cell and vascular panels remain at or below zero for every cluster. (C) Mean detection rate of each marker set per cluster, summarising the panel in (A). Vascular and guard-cell nuclei are therefore too rare in this dataset to form clusters of their own. (D) Expression of genes shared by all mesophyll clusters (shared core mesophyll genes) compared with genes specific to individual mesophyll clusters, scaled per gene across clusters. The shared core confirms a common mesophyll identity underlying Clusters 0, 1, 2, 4 and 5, while the cluster-specific sets show that these clusters differ as cell states rather than as cell types. (E) UMAP embedding split by treatment, with nuclei coloured by cluster identity and shaped according to whether they score higher on published palisade or spongy mesophyll marker sets (Xia et al. 2022). Palisade- and spongy-leaning nuclei are intermixed within every cluster and do not resolve into separate populations, reflecting the extensive overlap between the two published gene sets.

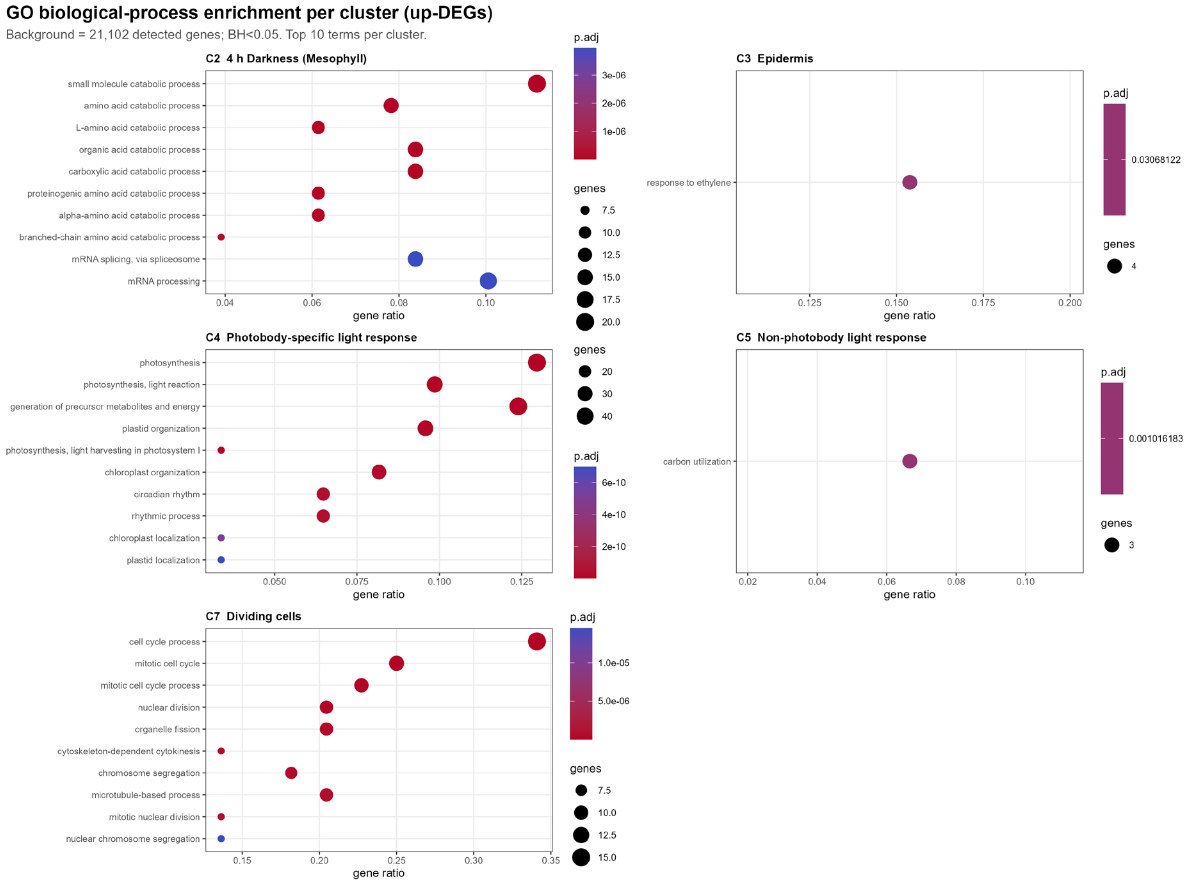


**Supplemental Figure 10: Gene Ontology (GO) term analysis of Clusters identified in snRNAseq dataset shows clear light-state signatures.** GO term ‘Biological Process’ analysis of Clusters from **Figure 7**. Circles depict # of genes per category, significance level (colour scale), and the ratio of the genes associated with the GO term vs the gene size of the whole Cluster.

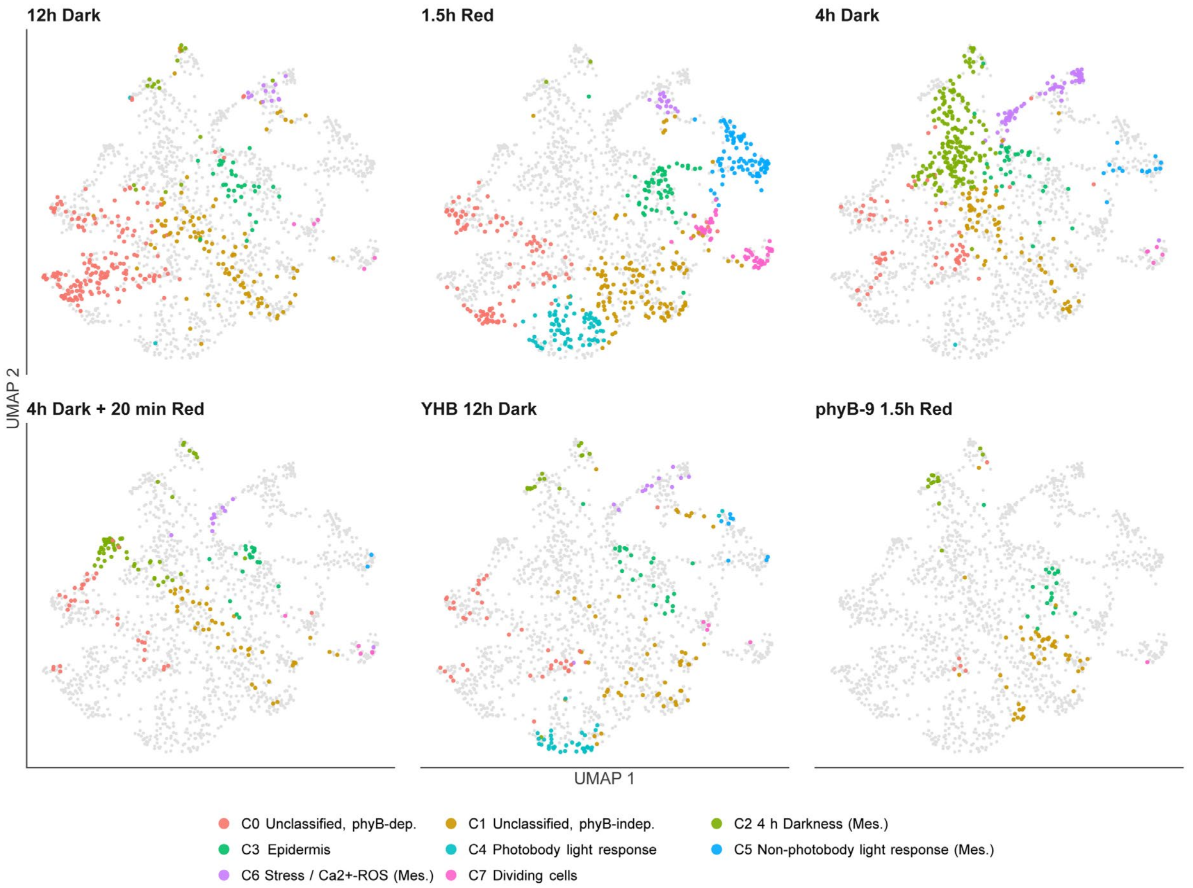


**Supplemental Figure 11: UMAP plots showing cluster enrichment per treatment.** UMAP plots accompanying Figure 7, showing cluster distribution per treatment.

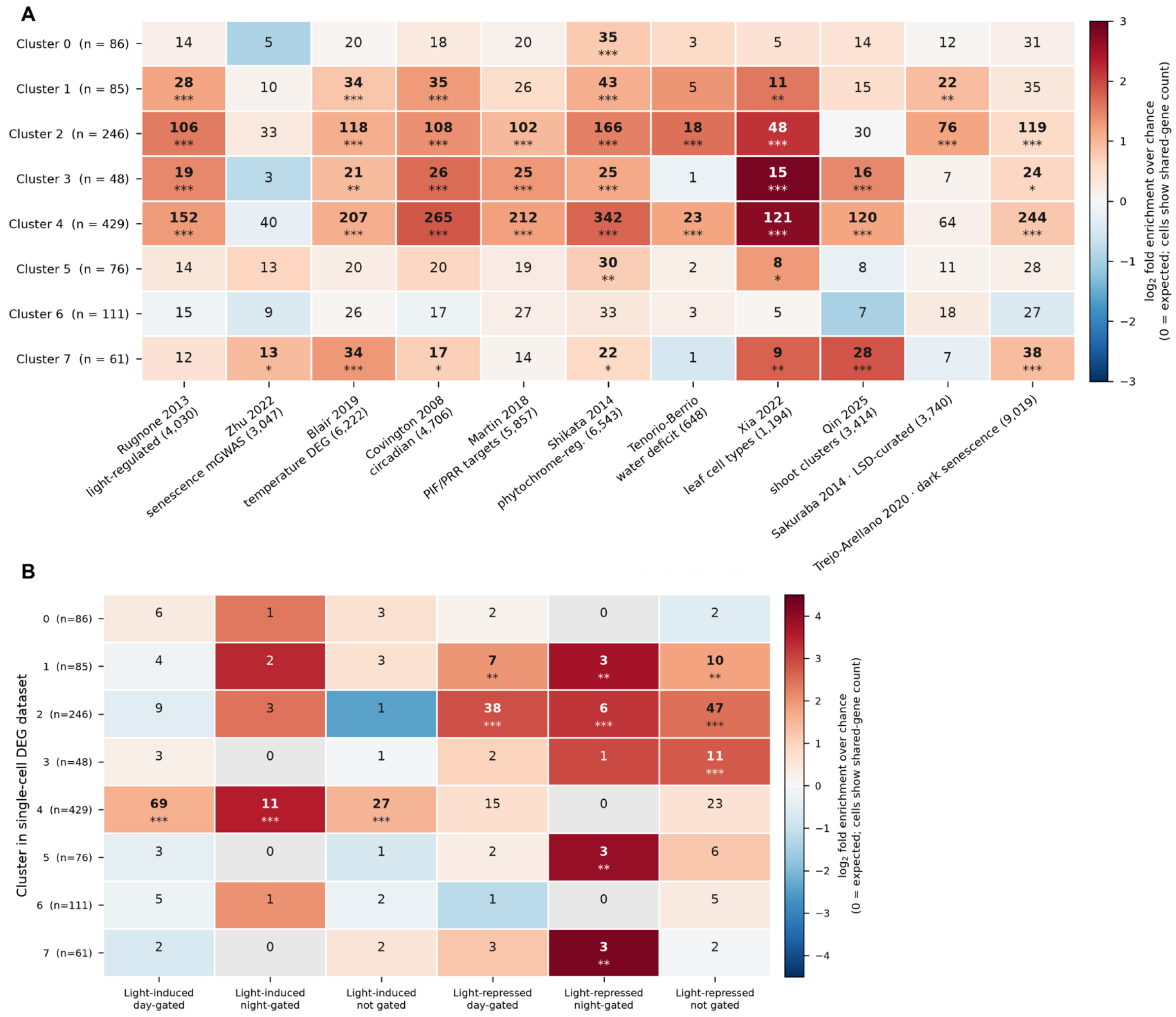

**Supplemental Figure 12: Statistically significant overlap between snRNA-seq dataset and existing transcriptomic datasets of light, circadian and senescence experiments. (A)** Number of genes per cluster that overlap with the whole DEG dataset indicated on the bottom of the columns. **(B)** Cluster-specific overlap for specific treatments in Rugnone et al. 2013. The colour scale shows the log2-fold enrichment of the particular cluster/dataset overlap. Asterisks denote statistically significant difference through Benjamini-Hochberg test for false-discovery rate, *p<0.05, **p<0.01, ***p<0.001.

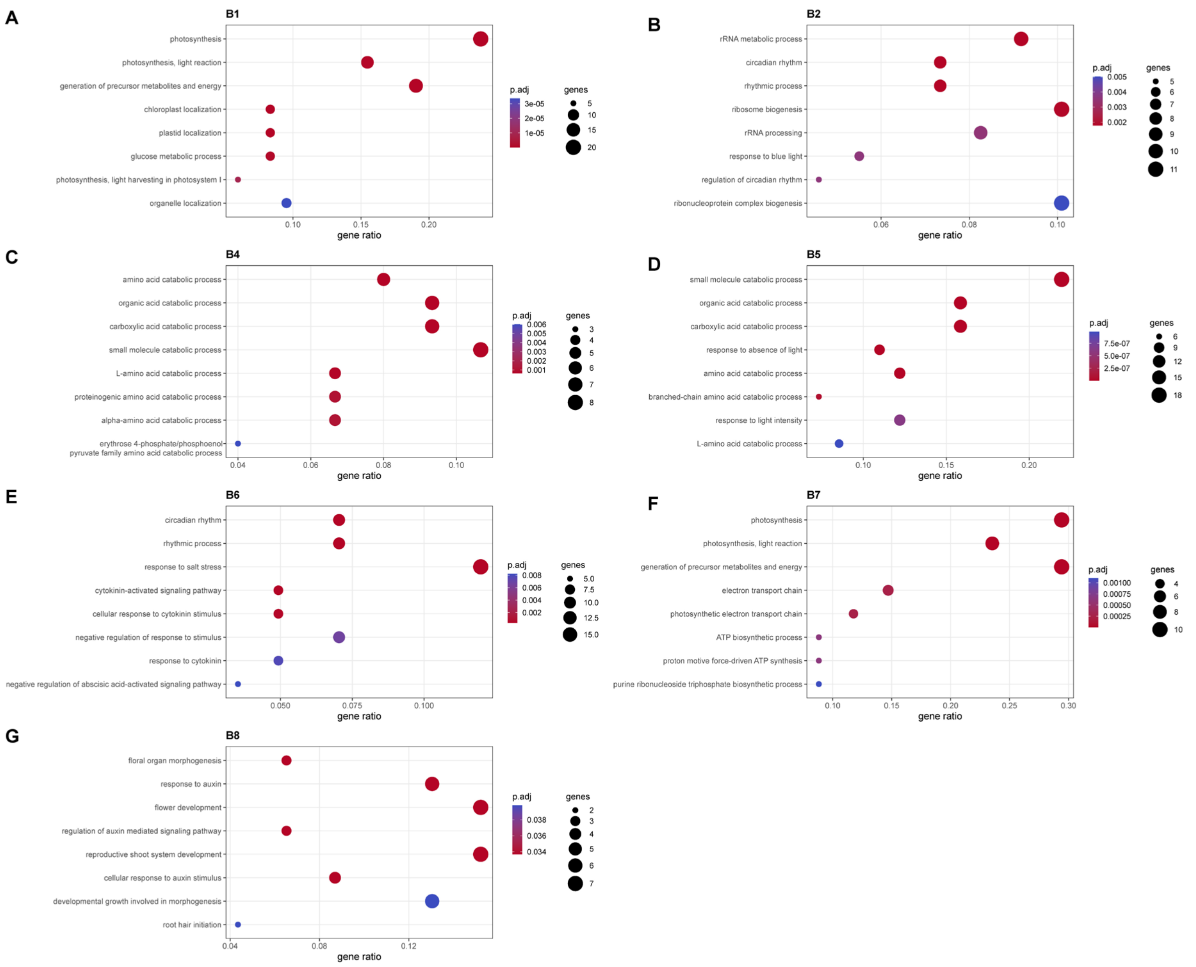


**Supplemental Figure 13. Gene Ontology (GO) term analysis of treatment and epidermis/mesophyll specific expression blocks.** GO term ‘Biological Process’ analysis of Blocks from **Figure 8** and **Supplemental file 5**. Circles depict # of genes per category, significance level (colour scale), and the ratio of the genes associated with the GO term vs the gene size of the whole expression Block.

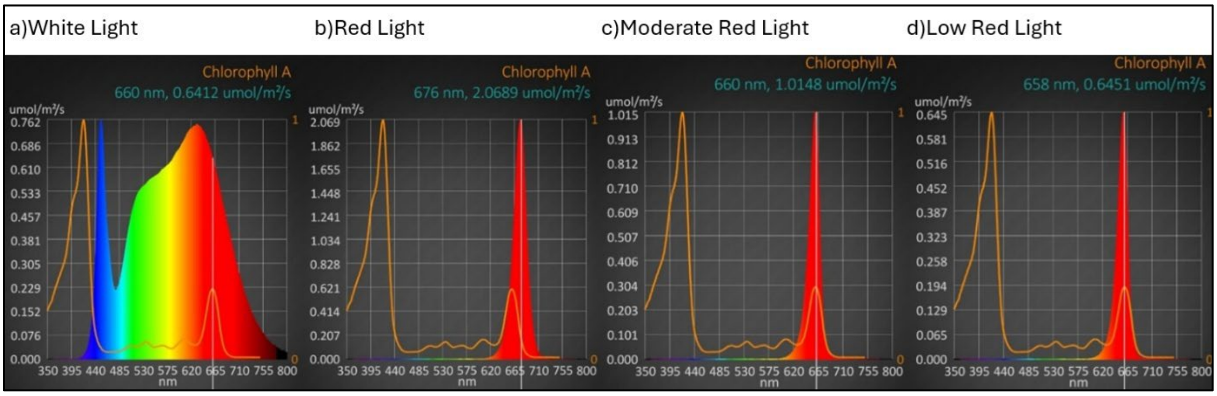


**Supplemental Figure 14. UPRtek spectrometer measurements of the applied light conditions.** Spectral profiles of (a) White Light, (b) Red Light, (c) Moderate Red Light, and (d) Low Red Light obtained with a UPRtek spectrometer. Measurements were recorded across 350–800 nm and are presented as photon flux density (μmol $m^{-2}$ $s^{-1}$) per wavelength.

| group | I_inf | A | k | t_half | absolute_t_half | I_half | mobile_fraction | immobile_fraction | n_files | n_outliers |
|---|---|---|---|---|---|---|---|---|---|---|
| Pavement Low-red | 68,02 | 55,97 | 0,12 | 5,77 | 6,77 | 40,04 | 0,64 | 0,36 | 23 | 1 |
| Pavement High-Red | 85,34 | 87,12 | 0,16 | 4,3 | 5,3 | 41,78 | 0,86 | 0,14 | 7 | 0 |
| WL-guard | 36,58 | 36,58 | 0,08 | 9,2 | 10,2 | 18,29 | 0,37 | 0,63 | 20 | 0 |
| WL-pavement | 51,03 | 51,03 | 0,15 | 4,69 | 5,69 | 25,51 | 0,51 | 0,49 | 19 | 0 |
| WL-palisade | 89,25 | 89,25 | 0,2 | 3,5 | 4,5 | 44,63 | 0,89 | 0,11 | 8 | 0 |
| WL-spongy | 72,28 | 72,28 | 0,21 | 3,26 | 4,26 | 36,14 | 0,72 | 0,28 | 9 | 0 |
| Guard Low-red | 33,41 | 32,73 | 0,07 | 10,35 | 11,35 | 17,04 | 0,33 | 0,67 | 29 | 0 |
| Guard High-Red | 30,98 | 31,37 | 0,14 | 5,07 | 6,07 | 15,29 | 0,31 | 0,69 | 15 | 1 |

**Supplemental Table 1. DynaFRAP output paramaters.** Parameters of FRAP data from Figure 4. (I_inf, A and k) Parameters of fitted curve. used to fit curves for FRAP data from Figure 4

| Symbol | AGI | log2FC | pct_in | pct_rest | adj_p | jung_list |
|---|---|---|---|---|---|---|
| LNK1 | AT5G64170 | 2.27 | 35 | 8 | 1.10E-16 | 17C |
| LNK3 | AT3G12320 | 2.1 | 28 | 6 | 1.40E-14 | 17C |
| phyB | AT2G18790 | 2.05 | 25 | 4 | 2.30E-18 | 17C, 27C |
| CAB4 | AT3G47470 | 1.76 | 52 | 21 | 7.00E-13 | 17C |
| MGP3 | AT1G68990 | 1.6 | 38 | 9 | 3.40E-18 | 17C |
| ATPIN3 | AT1G70940 | 1.22 | 31 | 10 | 2.80E-06 | 17C |
| DAR1 | AT4G36860 | 0.82 | 30 | 11 | 6.90E-04 | 17C |
| PIF5 | AT3G59060 | 0.78 | 46 | 22 | 1.20E-03 | 17C |

**Supplemental Table 2. Overlapping genes between Cluster 4 and Jung 2016 ChIP-seq phyB targets.**

**phyB photobodies display molecular memory, regulating light signaling transcriptional states through phase separation.**

### Supplemental methods

#### DynaFRAP analysis

Deriving recovery curves from the recorded Z-stack photobleaching time series requires tracking the bleached photobody and correcting its intensity for the acquisition-inherent signal fall-off using a non-bleached reference photobody. To this end, we employed a minimal Biom3d model, trained on ten manually annotated photobody sequences, to generate a 3D log-likelihood heatmap for every time frame, providing a voxel-wise proxy for the probability that a photobody is present. We then implemented an interactive viewer for exploring these heatmap sequences. For each time series, the user delineates bounding boxes in the pre-bleach frame that enclose the target (bleached) and the control photobody. Within each box, the position of the log-likelihood maximum is taken as the photobody centre. In the subsequent frame, the bounding box is re-centred on the previously detected centre, and the next centre is sought within this shifted subvolume. Centre detection is therefore propagated purely locally, independently of global network activations and thresholds. The mechanism consequently relies only on the sensitivity of the network to local structure, which proved sufficient in the majority of our recordings. Around each identified centre, geometric primitives are parametrised to sample and aggregate the image intensities. Our implementation circumscribes cylinders of fixed height (9 Z-slices) whose radius is estimated, separately for each time series, to contain at least 80% of the predicted photobody signal around the pre-bleach target centre. At any time frame, the user may adjust the bounding boxes and propagate the correction to all subsequent frames; after each propagation, a CSV file containing all results and the data required for full reproduction is exported and updated. Finally, the recorded intensities were corrected for photobleaching using the control photobodies, averaged across the groups of interest, and fitted with a one-phase exponential recovery model to extract features such as the half-time of recovery and the mobile fraction.

The processing pipeline is implemented as four modules: (a) pre-processing of LIF-format recordings into individual TIF-format time frames; (b) segmentation of the individual TIF files using the Biom3d module; (c) interactive measurement based on the segmentation results; and (d) post-processing of the measured positions and intensities to obtain the FRAP features and generate the corresponding charts. This modular architecture allows intermediate results to be reproduced independently and experiments to be adapted through configuration files alone, with dependencies managed by the uv package manager. The segmentation step requires a pre-trained Biom3d model checkpoint.

The proposed semi-automatic photobody tracking tool has the potential to accelerate the annotation process substantially. Tracking the centres in the nuclei of the comparatively immobile guard cells consistently required only a single propagation. The more granular pavement cells only rarely required one to two corrections, mainly in the later frames, where the photobody signal inherently degrades. Spongy and palisade mesophyll cells were the most demanding, with a few cases requiring corrections in nearly every frame, largely owing to the highly dynamic nature of their photobodies.

**Biom3D model training, pre- and post-processing code.**

### Training of the convolutional neural network and image pre-processing

A three-dimensional convolutional neural network was trained in Biom3D (Mougeot et al. 2024) using ten 3D image stacks and their corresponding manually annotated binary masks. Each stack was acquired as a 10-plane z-series and stored in TIFF format. The parameters extracted from the segmented objects were photobody number, photobody volume and mean photobody intensity. Inference in Biom3D requires the ImageJ-specific metadata header, which is not present in the raw acquisition files. All stacks were therefore re-saved through ImageJ before being passed to the trained model. This conversion step was automated with a Python script (Figure 1).

```
import subprocess
import glob
import os

#folder paths
input_folder = r"C:\Users\AGKvG\Desktop\Protocol For Automated Image Annotation and Analysis\raw"
output_folder = r
"C:\Users\AGKvG\Desktop\Protocol For Automated Image Annotation and Analysis\raw images processed in ImageJ"
macro_path = r"C:\Users\AGKvG\ImageJ macro code.ijm"

#path to your ImageJ or Fiji executable
imagej_path = r"C:\fiji-win64\Fiji.app\ImageJ-win64.exe"

#find all TIFF files in the input folder
tif_files = glob.glob(os.path.join(input_folder, "*.tif"))

#loop through each TIFF file and process it with ImageJ
for tif_file in tif_files:
    filename = os.path.basename(tif_file)
    output_file_path = os.path.join(output_folder, filename)  # Full path including filename for output

    #construct the command to run ImageJ with macro and arguments
    command = f'"{imagej_path}" --headless -macro "{macro_path}" "{tif_file},{output_file_path}"'

    # Run the command
    subprocess.run(command, shell=True)
```

Figure 1. Code for ImageJ image saving automatization.

The script defined the input directory, the output directory, the path to an ImageJ macro containing the conversion instructions and the path to the ImageJ executable. All TIFF files in the input directory were collected with the glob module, and each file was processed in turn: the output path was constructed from the output directory and the original filename, so that file names were preserved between input and output. ImageJ was invoked in headless mode through the subprocess module, with the macro and the input and output paths supplied as command-line arguments, allowing the full image set to be converted without user interaction. The trained model was then applied to the converted stacks to generate binary masks for all remaining images.

### Automated 3D image analysis of multi-channel TIFF files

Segmented objects were quantified with a custom Python script (Figure 2) that operated on two inputs per sample: the binary mask produced by the trained model, which defines the objects of interest, and the corresponding raw stack, which provides the voxel intensity values. The script used numpy for array handling, tifffile for image input and output, scipy.ndimage for connected-component labelling, skimage.measure for region property measurement, and pandas for tabulation.

Analysis was implemented in a function (process_image_analysis) taking the mask path, the raw data path and the output path of the results table as arguments. After both files were loaded, the array dimensions were inspected to determine the number of channel or time slices (T) and the number of z-planes (Z). The total number of planes was verified to be an integer multiple of the number of T slices, and the mask array was reshaped accordingly to separate the T and Z dimensions. Connected components were then labelled in three dimensions for each T slice independently, using scipy.ndimage.label with a defined 3D structuring element, so that voxels belonging to a single photobody across consecutive z-planes were assigned a common label. For every labelled region, skimage.measure.regionprops was applied to the corresponding raw intensity data to extract object volume (voxel count), mean intensity and integrated intensity. The measurements for all objects were collected into a pandas DataFrame and written to a CSV file.

The function was applied in batch. The script iterated over all mask files in the mask directory, derived the name of the matching raw data file from a fixed naming convention, and verified its presence before analysis. Samples for which no matching raw file was found were skipped and reported to the console, and the output directory was created automatically if it did not already exist.

```
#USE THIS CODE SNIPPLET ONLY FOR TIFF FILES WHICH HAVE BOTH Z AND T DIMENSION

import os
import numpy as np
import tifffile
from scipy.ndimage import label
from skimage.measure import regionprops
import pandas as pd

def process_image_analysis(mask_path, raw_path, output_csv_path):
    # Load binary mask and raw data
    binary_mask = tifffile.imread(mask_path)
    raw_data = tifffile.imread(raw_path)

    # Calculate T and Z channels for reshaping
    t_channels_raw = raw_data.shape[0]
    z_slices_per_t_channel = binary_mask.shape[0] // t_channels_raw

    # Validate division
    assert binary_mask.shape[0] % t_channels_raw == 0, "Mismatch in division between T channels and total slices"

    # Reshape binary mask to T and Z channels
    reshaped_mask = binary_mask.reshape((t_channels_raw, z_slices_per_t_channel, binary_mask.shape[1], binary_mask.shape[2]))

    # Label connected components with 3D connectivity
    labeled_stacks = np.zeros_like(reshaped_mask, dtype=int)
    for i in range(reshaped_mask.shape[0]):
        structure = np.ones((3, 3, 3))
        labeled_stacks[i], _ = label(reshaped_mask[i], structure=structure)

    # Initialize list for measurements
    measurements = []

    # Loop through T channels for metrics calculation
```

**Figure 2.** Python script for automated 3D object labelling and measurement in multi-channel TIFF stacks.

#### Standardisation of measurement tables

The per-sample measurement tables were reformatted with a further Python script (Figure 3) to match the input specification of the downstream R plotting pipeline. The script used pandas for data handling, os for file system operations and re for regular-expression matching, and created the output directory if it was absent.

All files in the input directory whose names ended in _measurements.csv were processed. Experimental metadata (for example acquisition date and experiment type) were parsed from each file name using a regular-expression pattern. Each table was then read into a DataFrame and a Filename column was added, containing the file name with the _measurements.csv suffix removed, so that every measurement could be traced to its source image after the tables were pooled. The columns T_Channel and Label were renamed Timeslot and PhotobodyID, respectively. The processed tables were written to the output directory as CSV files without the index column. File names that did not match the expected pattern were reported to the console and left unprocessed.

```python
import pandas as pd
import os
import re

def process_csv_files(input_dir, output_dir):
    # Compile the regex pattern for extracting information from the filenames
    pattern = re.compile(
        r'(\d{2}-\d{2}-\d{2})\s+([A-Z]+).*?(\d+)min(\d+)sec\d+\s*(R|FR).*?_measurements.csv'
    )

    # Ensure output directory exists
    os.makedirs(output_dir, exist_ok=True)

    # Iterate over all files in the input directory
    for filename in os.listdir(input_dir):
        if filename.endswith('_measurements.csv'):
            # Extract information from the filename
            match = pattern.match(filename)
            if match:
                # Removed unnecessary variables since we're not including them in the dataframe

                file_path = os.path.join(input_dir, filename)
                df = pd.read_csv(file_path)

                # Add the Filename column without unnecessary ones
                df['Filename'] = filename.replace('_measurements.csv', '')

                # Rename the T_Channel to Timeslot and Label to PhotobodyID
                df.rename(columns={'T_Channel': 'Timeslot', 'Label': 'PhotobodyID'}, inplace=True)

                # Before saving, ensure we do not include the previously mentioned columns by not adding them at all

                # Save the modified dataframe
                output_file_path = os.path.join(output_dir, filename)
                df.to_csv(output_file_path, index=False)
                print(f"Processed {filename}")
            else:
```

**Figure 3.** Python script for renaming and reformatting the measurement tables.
